# High-efficiency likelihood inference of shared proteomic architectures across 50 complex human traits

**DOI:** 10.1101/2025.03.16.643065

**Authors:** Xiaoru Sun, Sizhe Yang, Qianqian Peng, Xuan Zhang, Yijia Qian, Renliang Sun, Guoqing Zhang, Sijia Wang, Li Jin, Menghan Zhang

## Abstract

Advancements in genetic correlation estimation have elucidated genome-wide pleiotropy’s influence on phenotypic correlations among human complex traits and diseases. However, the role of proteomic domains in these correlations remains underexplored. Traditional genetic correlation analysis assumptions, including the minute effects of SNPs and their linkage disequilibrium, do not suit proteomic data. We present a novel method, Likelihood-based Estimation for Proteomic Correlation (LEAP), tailored to provide unbiased estimation of shared proteomic architectures between trait pairs. LEAP notably decreases computational demands by approximately 1000-fold compared to conventional bivariate linear mixed models. We applied LEAP to data from the UK Biobank Pharma Proteomics Project, identifying 585 significant proteomic correlations among 1,225 pairs of 50 biochemical, anthropometric, and behavioral traits. Furthermore, we quantified the distinct proteomic and genetic contributions to phenotypic correlations, highlighting significant gender differences. This study provides a comprehensive computational approach for proteomic correlation estimation, clarifying the specific roles of genomics and proteomics in complex trait correlations. Our findings not only advance the understanding of proteomic contributions to phenotypic traits but also suggest potential applications for evaluating shared omics architectures in other domains such as transcriptomics and metabolomics.

## Background

The pleiotropic effects observed across various levels of omics significantly influence the plasticity of phenotypic interactions in humans[1, 2], thereby inducing correlations among human phenotypes[3]. Analyzing these phenotypic correlations can illuminate the underlying genetic architectures, elucidate gene functions and disease mechanisms, and offer novel insights for therapeutic innovations across a spectrum of human diseases[4]. Given appropriate omics data, it is possible to decompose phenotypic correlations into distinct omics components by estimating relevant measures, such as genetic correlation[5].

Recent advances in estimating genetic correlation have provided valuable insights into the phenotypic correlations between traits induced by genome-wide pleiotropy[5-7]. However, human genetics alone is insufficient for fully understanding the biology or mechanisms underlying the formation of complex traits and phenotypic interactions, as genome-wide association studies (GWAS) frequently identify genetic variants without pinpointing causal genes mediating their impacts[8]. The development of high-throughput proteomics provides new insights into proteomic signatures associated with human traits[9], since proteins are the key performers in expressing genetic information and essential functional units of the human body[10]. Proteomics is therefore crucial for investigating the genetic mechanisms of complex traits, revealing specific biological mechanisms of the intermediate cellular molecular processes, and potentially uncovering the joint effects of genetics and environmental exposures on complex traits[11].

Integrating proteomics with genomic and phenotypic information may bridge the gap between the human genome and human traits[12]. However, there is a limited overlap between GWAS signals and protein quantitative trait loci (pQTL)[13-15]. This discrepancy may be partly due to insufficient sample sizes in proteomic or genetic data, resulting in lower pQTL detection capabilities, especially in tissues other than blood. Additionally, a significant portion of proteomic signals cannot be explained by GWAS signals alone, suggesting that proteomics may involve both genetic transmission effects and proteomic-specific effects on complex traits[16-18]. Thus, it is necessary to develop proteomics-based approaches to systematically analyze the impact of the proteomic layer on the formation and interactions of complex traits.

Despite the continuous increase in proteomic data, there is still a lack of research on the contributions of proteomic domains beyond the genome to the plasticity of phenotypic interactions. Exploring the shared proteomic architecture between traits, known as the proteomic correlation, remains challenging. One prominent reason is that prevailing computational approaches are primarily designed for estimating genetic correlation[7, 19-22], such as the cross-trait Linkage Disequilibrium Score Regression model (LDSC)[7] and the high-definition likelihood model (HDL)[21]. Notably, there are three incompatibilities for utilizing genetic correlation approaches to estimate proteomic correlations. First, the architectures of proteomic data exhibit total differences from genetic ones. When estimating genetic correlations, SNPs are usually assumed to be independent across long-distance or on different chromosomes[21]. Proteins from different genomic loci or chromosomes can still exhibit high correlations[23]. Second, the physical positions of SNPs in the genome are fixed[24], which contrasts with the dynamic nature of proteins[25]. Third, most genetic effects on complex traits are infinitesimal[26], whereas proteomic effects are arbitrary. Ignoring these differences can lead to biased estimates of proteomic correlation. Consequently, computational approaches for genetic correlation estimation cannot be directly applied to proteomic data. In addition, the bivariate linear mixed model (LMM), which relies on random effects, can capture shared proteomic architectures between correlated traits[27, 28]. However, standard fitting algorithms for LMM, including the traditional restricted maximum likelihood estimation method (REML), exhibits limited computational efficiency on large-scale data[29, 30]. Therefore, it is essential to develop a general computational approach for proteomic correlation estimation, considering the properties of proteomic data while achieving high efficiency.

Given these considerations, we here presented a <u>L</u>ikelihood-based method for <u>E</u>stim<u>A</u>ting the <u>P</u>roteomic Correlation (LEAP) between traits (Fig. 1). The rationale behind LEAP is to estimate the variance/covariance components based on marginal association test statistics (i.e., the summary statistics from proteome-wide association studies) after correcting for covariance matrixes of the proteomic data. It is noteworthy that our approach relaxes computational constraints derived from primary genetic architectures to enhance applicability to proteomic data (Table 1). We assessed the accuracy and efficiency of LEAP compared with LMM through simulations. In the empirical applications, we estimated the proteomic correlations among 50 complex traits encompassing biochemical, anthropometric, and behavioral measurements for 43,509 White British samples in the UK Biobank (UKBB). We also examined the gender differences in these empirical applications and explained the statistical findings using genome-wide pQTL analysis. Our work illustrates the computational benefits of LEAP and highlights the distinct roles of genomics and proteomics in the plasticity of phenotypic interactions.

**Table 1.**
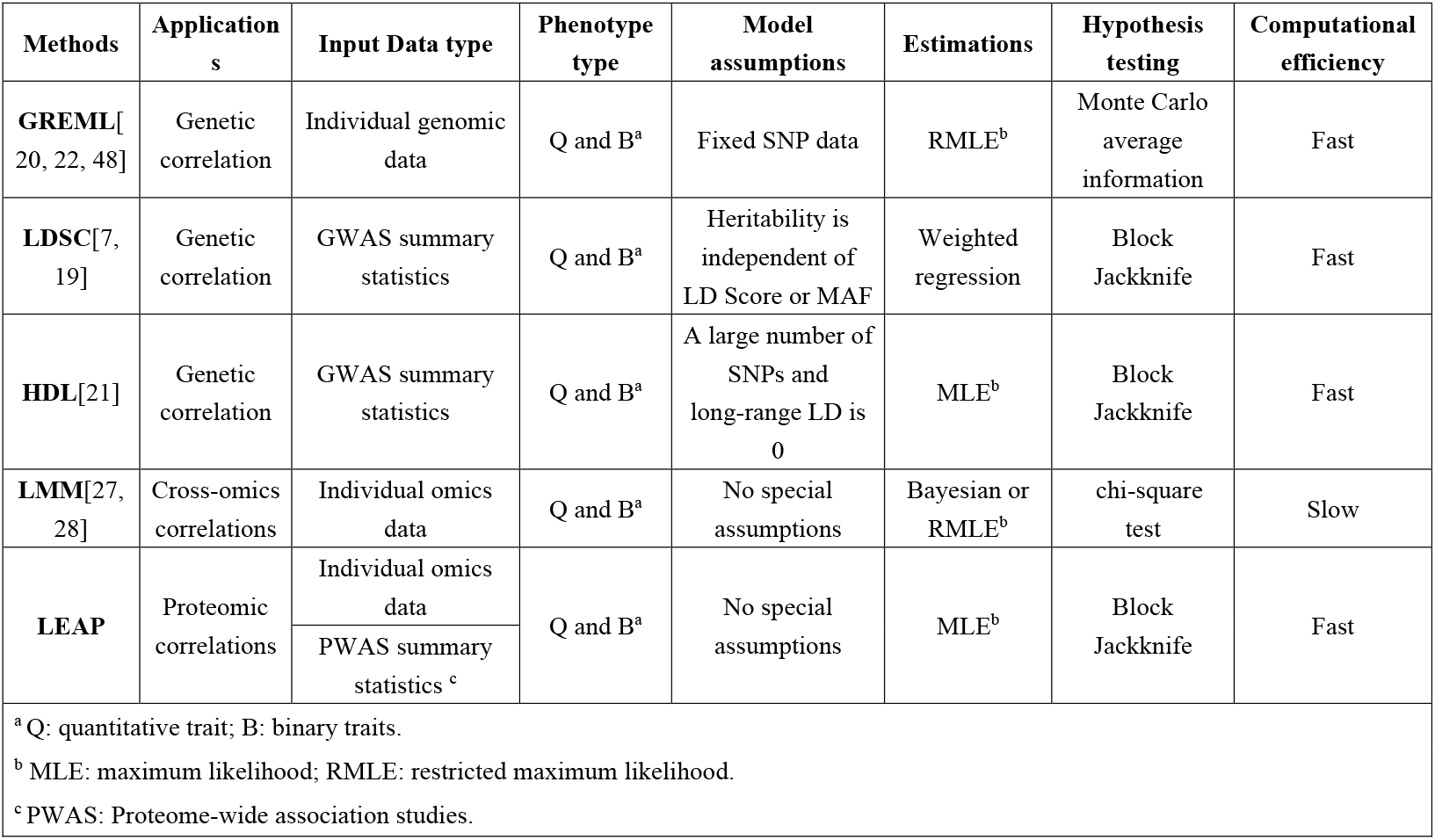
Summary of some existing phenotypic correlation estimation methods.

| Methods | Applications | Input Data type | Phenotype type | Model assumptions | Estimations | Hypothesis testing | Computational efficiency |
| --- | --- | --- | --- | --- | --- | --- | --- |
| **GREML**[20, 22, 48] | Genetic correlation | Individual genomic data | Q and B<sup>a</sup> | Fixed SNP data | RMLE<sup>b</sup> | Monte Carlo average information | Fast |
| **LDSC**[7, 19] | Genetic correlation | GWAS summary statistics | Q and B<sup>a</sup> | Heritability is independent of LD Score or MAF | Weighted regression | Block Jackknife | Fast |
| **HDL**[21] | Genetic correlation | GWAS summary statistics | Q and B<sup>a</sup> | A large number of SNPs and long-range LD is 0 | MLE<sup>b</sup> | Block Jackknife | Fast |
| **LMM**[27, 28] | Cross-omics correlations | Individual omics data | Q and B<sup>a</sup> | No special assumptions | Bayesian or RMLE<sup>b</sup> | chi-square test | Slow |
| **LEAP** | Proteomic correlations | Individual omics data | Q and B<sup>a</sup> | No special assumptions | MLE<sup>b</sup> | Block Jackknife | Fast |
|  |  | PWAS summary statistics <sup>c</sup> |  |  |  |  |  |
| <sup>a</sup> Q: quantitative trait; B: binary traits. |  |  |  |  |  |  |  |
| <sup>b</sup> MLE: maximum likelihood; RMLE: restricted maximum likelihood. |  |  |  |  |  |  |  |
| <sup>c</sup> PWAS: Proteome-wide association studies. |  |  |  |  |  |  |  |

**Fig. 1.**
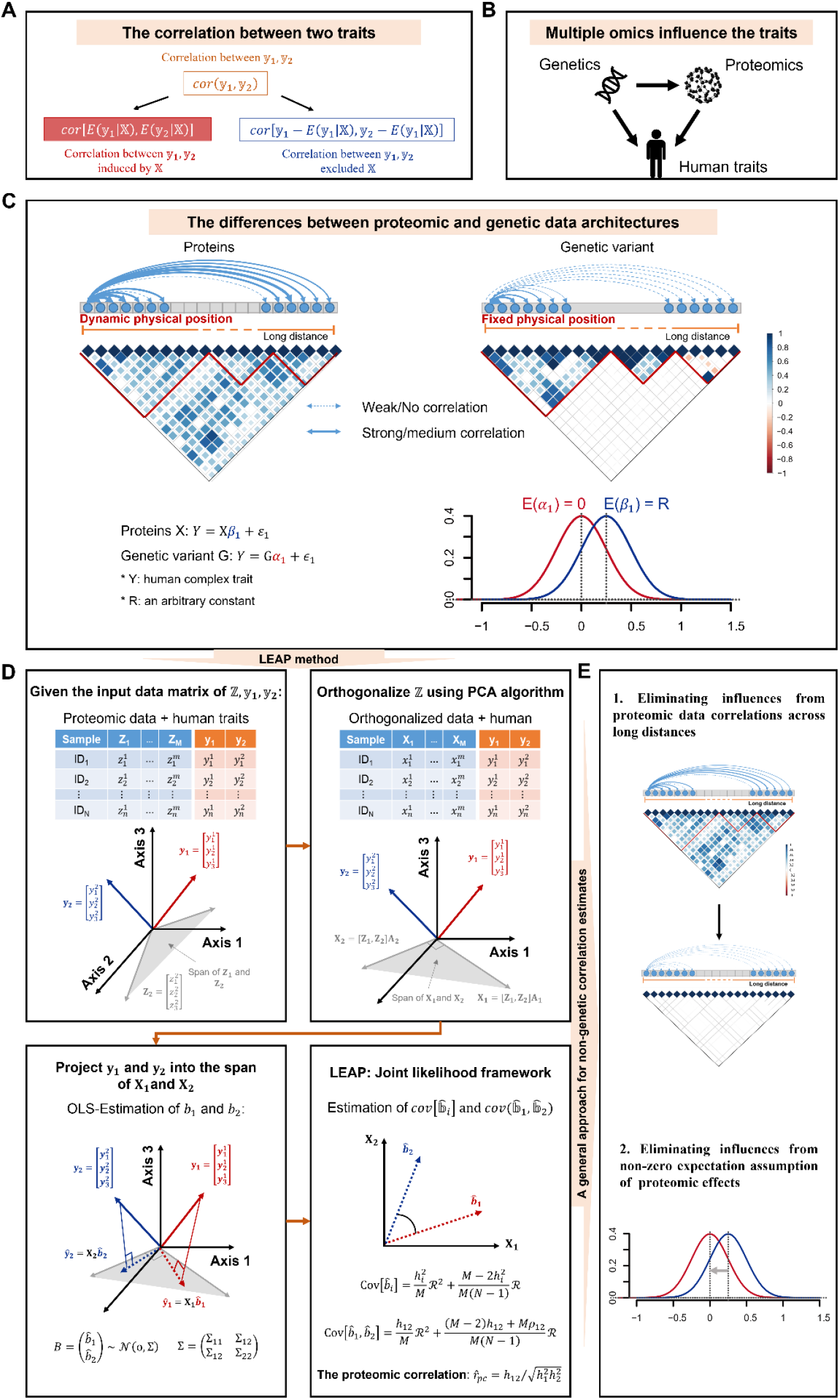
Overview of the LEAP method. (A) The phenotypic correlation between two traits can be divided into two components: the correlation induced by the shared omics data (**X**) and the residual correlation excluding **X**. (B) Human complex traits are influenced by multiple omics, such as genetics and proteomics. (C) Differences in data architectures between genetic variants and proteins, including the physical positions and proteomic correlations, as well as the effect sizes of genetic variants and proteins on traits. (D) The workflow of the individual-level data version of LEAP estimates the proteomic correlation between two traits. Considering the proteomic architectures, we performed a linear orthogonal transformation on proteomic data using the PCA algorithm to eliminate the influences from long-distance proteomic data correlations. Based on the relationship between the observed proteomic effects on two traits, the true proteomic correlation can be obtained using log-likelihood estimations. (F) LEAP eliminates the incompatibilities from non-genetic data and reports the non-genetic correlations between pairwise human traits.

## Results

### Overview of the LEAP method

LEAP is designed to estimate the proteomic correlations between two traits via aggregating the effects of proteins (i.e., proteomic effects) on the traits. Suppose there is a cohort for two traits with sample sizes *N. Z =* {*Z*_*j*_, *j =* 1, …, *M*} denotes a *N* × *M* matrix of the proteomic data, where *M* represents the number of proteins. *X* is a *N* × *M* matrix of the proteomic data after the linear orthogonal transformation, i.e., *X = ZV*, where *V* is the right singular matrix of *Z*(*Z = UΛ*_1_*V*^*T*^). Without loss of generality, we assume that *X* is scaled to be zero mean and unit variance. The confounding factors are absent and two quantitative traits *y*_1_ and *y*_2_ are tested. The 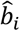 -vector denotes the estimated marginal effects from *X* to *y*_*i*_. Then the covariance matrices of 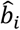 are given as

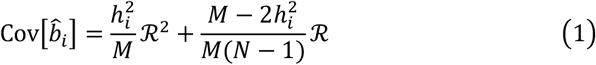

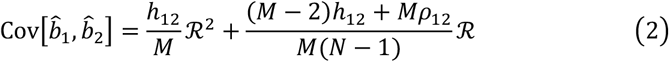

where ℛ is the *M* × *M* covariance matrix of the proteomic data, and 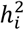 is the proportion of phenotypic variation due to the additive proteins we are interested in, which is defined as the proteomic explainability. *h*_12_ is the proteomic covariance between the two traits. *ρ*_12_ represents the covariance of residuals between the two traits. Let

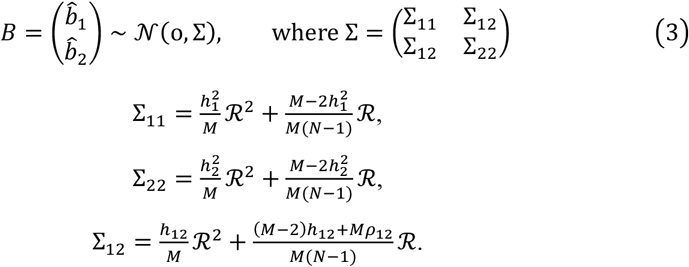

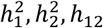 can be estimated by maximizing the full joint log-likelihood function,

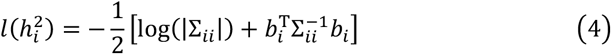

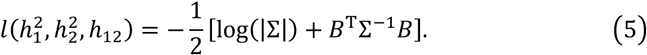

Then the proteomic correlation can be obtained as 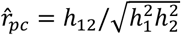.

Additionally, LEAP can also be applied to the summary-level data if the summary statistics (the marginal regression effect *μ* and its standard error) from PWAS and the proteomic covariance matrix (*Z*^*T*^*Z*) are available. The eigenvalue of ℛ (ℛ *= Q*Λ*Q*^*T*^) can be obtained from the eigen-decomposition of the proteomic covariance matrix, *Z*^*T*^*Z*/(*N* − 1) *=* VΛ*V*^*T*^. The orthogonal matrix *Q = V*^*T*^*V*. The transformed marginal effect of *X* on *y*_*i*_ can be obtained

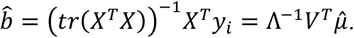

Then, the proteomic correlation can be estimated using the summary statistics as the same as using the individual-level data. We derived the above equations in Supplementary Note 1.1-1.5.

### Simulation results

Based on a series of simulation tests, we assessed LEAP’s accuracy, statistical precision, and computational efficiency under varying conditions, including sample size, the number of simulated proteins, and true proteomic correlation, comparing it to the LMM method. Considering the expensive computational burden of LMM in large-scale datasets, we utilized the full eigen decomposition of the simulated protein data instead of the raw *N*×*N* data covariance matrix for LMM fitting (i.e., the modified LMM, details can be found in Method and Supplementary Note). Initially, we simulated a relatively smaller dataset to test the performance of LEAP, the raw LMM and the modified LMM. Subsequently, we compared LEAP with the modified LMM with moderate sample sizes and evaluated LEAP alone with larger sample sizes. We generated the proteomic data matrix from a scaled multivariate normal distribution. The true effect sizes of each protein on the phenotypes were drawn from a bivariate normal distribution. The quantitative phenotypic traits were generated by adding errors from another bivariate normal distribution.

### Performance of LEAP, raw LMM and modified LMM with small sample sizes

We conducted simulations involving two traits and proteomic data under varying conditions, including sample size (N = 1,000, 2,000, 3,000, 4,000 and 5,000), the number of simulated variables (M = 50, 100, 150, 200, and 300) and true proteomic correlations between two traits (*r*_*pc*_ ranging from -0.9 to 0.9). The trait variance explained by proteomics (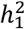 and 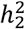) was set at 0.8 for both traits. All three methods obtained stable and unbiased estimates of the proteomic correlations with different settings (Fig. 2A). Notably, the statistical precision of all the three methods improved as the number of simulated variables and the absolute magnitude of true proteomic correlations increased.

**Fig. 2.**
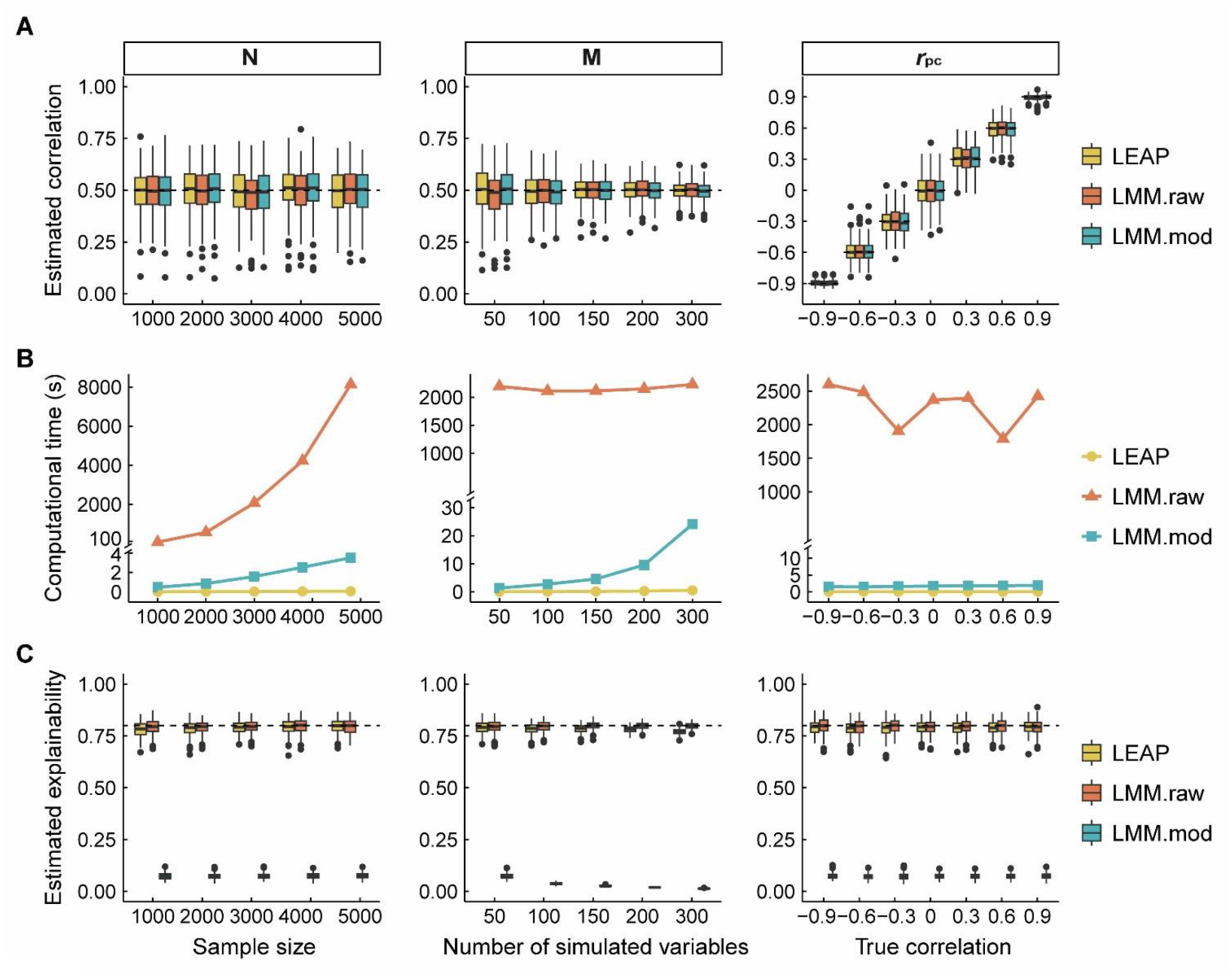
Simulation performance of LEAP against LMM. Both the raw LMM and the LMM with PC-based regularizations as inputs were tested. (A) The box plots of estimated proteomic correlations under varying conditions for the three methods: LEAP (yellow), raw LMM (orange) and modified LMM (blue). The sample sizes (N) were set at 1000, 2000, 3000, 4000 and 5000. The number of simulated variables (M) were set at 50, 100, 150, 200, and 300. The true correlation values (*r*_*pc*_) ranged from -0.9 to 0.9. 1000 replicated were simulated. (B) The computational time (in seconds) required by each method under the same varying conditions. (C) The box plots of estimated proteomic explainability for each method. The true proteomic explainability was set at 0.8 for both traits. Inside each box, the dash lines represent the true values, the solid lines indicate the median of the estimated values, the central boxes indicate the interquartile range (IQR), and the whiskers extend up to 1.5 times the IQR.

We further compared the computational time for each method under different conditions (Fig. 2B). LEAP exhibited significantly lower computational burdens than both the raw and modified LMM methods under identical parameter settings. As the sample size increased, the computational time of the raw LMM underwent a substantial increase, whereas LEAP remained relatively stable. Although the modified LMM exhibited higher computational efficiency than raw LMM, it was still less efficient than LEAP. As the number of simulated proteins increased, the computational efficiency of both LEAP and raw LMM decreased slightly, while that of modified LMM decreased significantly. Despite these variations, all the three methods maintained stability across different true correlation settings.

Additionally, we scrutinized the estimates of proteomic explainability using LEAP, raw LMM and modified LMM. Both LEAP and raw LMM demonstrated robustness in estimating proteomic explainability under varying conditions (Fig. 2C). However, the modified LMM might exhibit reduced ability to estimate proteomic explainability accurately.

### Performance of LEAP and modified LMM with moderate sample sizes

The performance assessment involved varied sample sizes (N = 1,000, 2,000, 3,000, 4,000, 5,000, 10,000, and 20,000), number of simulated proteins (M = 50, 100, 200, 500, and 1,000), and different true proteomic correlation settings (*r*_*pc*_ ranging from -0.9 to 0.9). In contrast, the raw LMM encountered calculation overflow under the same computational conditions, preventing a direct comparison with LEAP at this stage. We also evaluated proteomic explainability with configurations both high 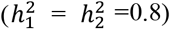, one high and one low (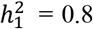 and 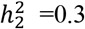), and both low 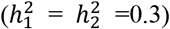 of the proteomic explainability (Fig. 3). The performance of LEAP was comparable to that of modified LMM and demonstrated enhanced accuracy in estimating ground-truth correlation coefficients (Fig. 3A to Fig. 3C). However, as the number of simulated proteins increased, LMM tended to underestimate correlations, particularly when the proteomic explainability was low for one or both traits (Fig. 3B).

**Fig. 3.**
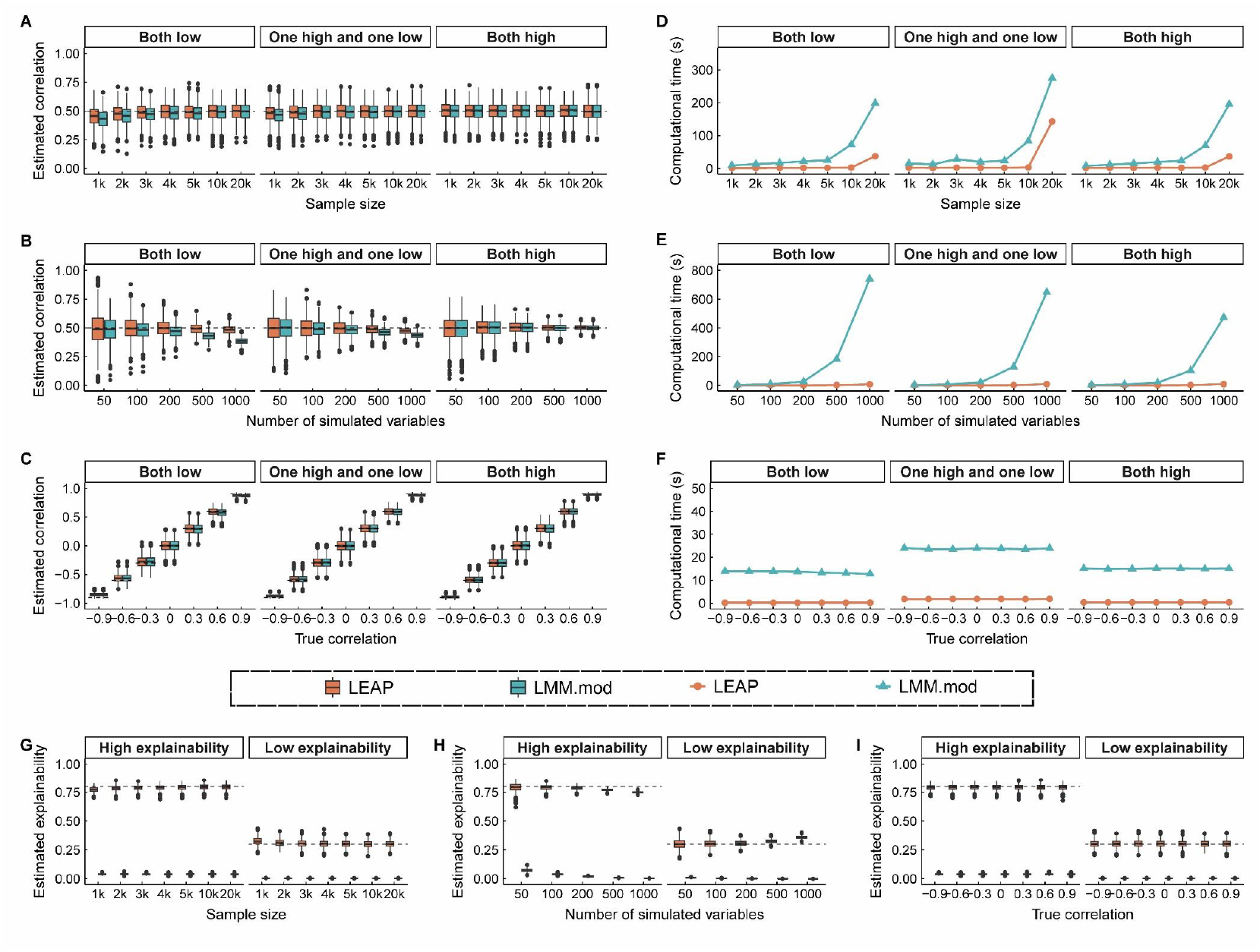
Simulation performance of LEAP against modified LMM under varying conditions. (A) shows the box plots of estimated proteomic correlations for LEAP (orange) and LMM (blue) under varying proteomic explainability (0.3 and 0.8) and sample size (1000, 2000, 3000, 4000, 5000, 10000, and 20000). Both low indicates both traits have low proteomic explainability (set at 0.3). One high and one low represent one trait has high explainability (set at 0.8), and the other has low explainability (set at 0.3). Both high indicates both traits have high proteomic explainability (set at 0.8). (B) shows the box plots of estimated correlations for varying numbers of simulated variables (50, 100, 200, 500, and 1000) under the same proteomic explainability conditions as in (A). (C) presents the box plots of estimated correlations against true correlation values (−0.9 to 0.9) under the same proteomic explainability conditions. Subfigures (D) to (F) illustrate the computational time (in seconds) required by each method across different sample sizes, numbers of simulated variables and true correlations under the same proteomic explainability conditions as in (A). Subfigures (G) to (I) display the box plots of estimated explainability for LEAP and LMM methods across different sample sizes, numbers of simulated variables and true correlations under high (set at 0.8) and low (set at 0.3) explainability conditions. Inside each box, the dash lines represent the true values, the solid lines indicate the median of the estimated values, the central boxes indicate the interquartile range (IQR), and the whiskers extend up to 1.5 times the IQR.

Notably, LEAP consistently exhibited significantly lower computational burdens compared to the modified LMM under identical parameter settings. As the sample size exceeded 5,000, the computational time for both LEAP and LMM saw substantial increases (Fig. 3D). However, LEAP’s computational efficiency remained stable despite increases in the number of simulated proteins and the magnitude of true correlations. In contrast, LMM’s computational time increased significantly with more simulated proteins (Fig. 3E and Fig. 3F).

For estimating proteomic explainability, LEAP consistently produced unbiased estimates, except in scenarios with a sample size of 5,000 and over 500 simulated proteins. In contrast, the modified LMM consistently exhibited a reduced ability to estimate proteomic explainability accurately across varying conditions (Fig. 3G to Fig. 3I).

### Efficiency of LEAP with larger sample sizes

We noted that the increased number of simulated proteins might bias with increased simulated proteins, especially when the estimations of LEAP when the ratio of sample size to the number of simulated variables (N/M) is less than 10. Therefore, we evaluated the accuracy and efficiency of LEAP under different ratios (N/M = 30 and N/M = 50) and larger sample sizes (N ranging from 10,000 to 200,000). Our results confirmed that LEAP consistently generated approximately unbiased estimations (Fig. 4A and Fig. 4B). Notably, higher proteomic explainability led to greater accuracy compared to lower values (Fig. 4B).

**Fig. 4.**
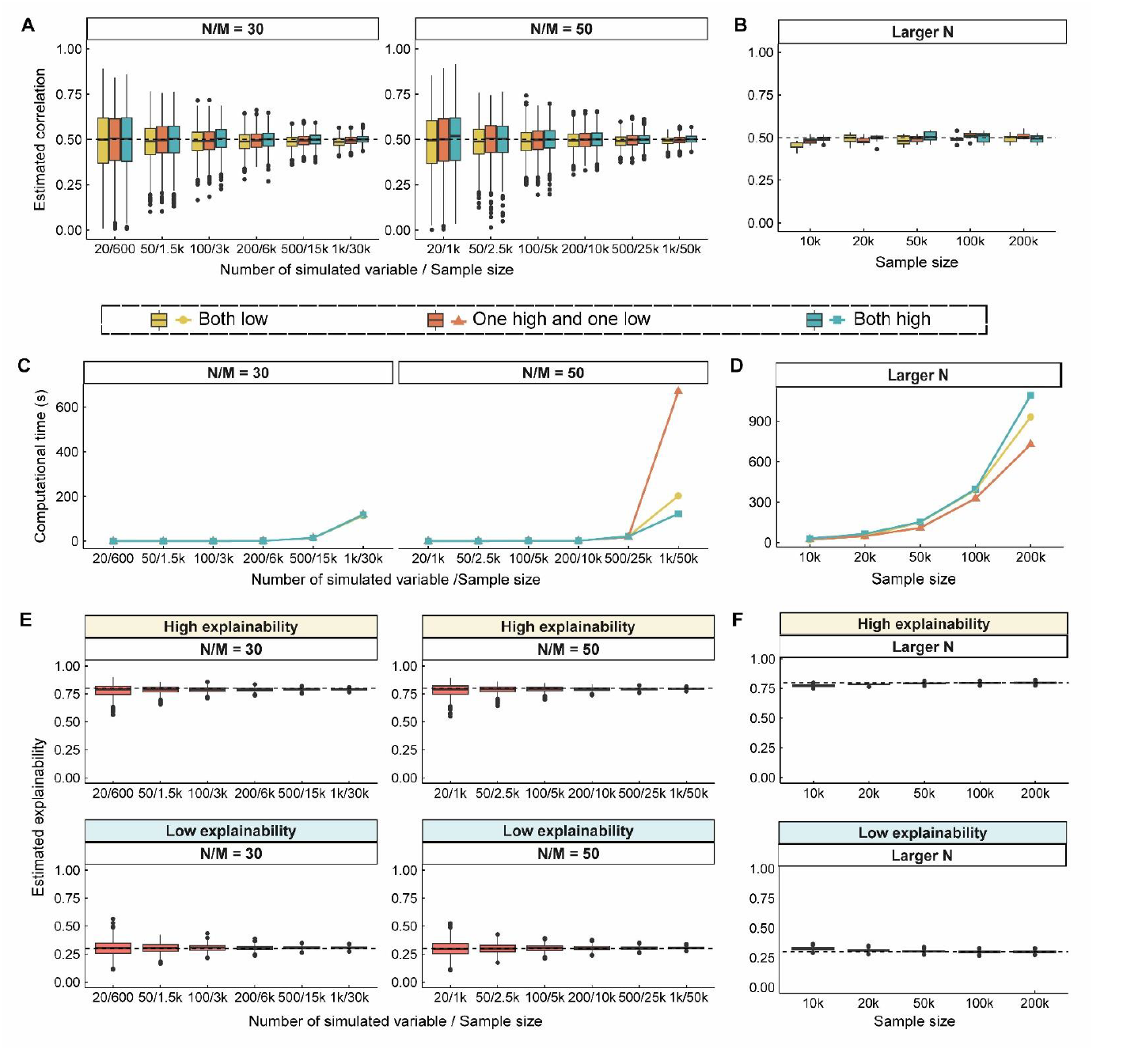
Simulation performance of LEAP under various conditions. (A) shows the box plots of estimated proteomic correlations for LEAP under different proteomic explainability conditions (both low in yellow, one high and one low in orange and both high in blue), and different ratios of the number of simulated variables to sample size (N/M = 30 and N/M = 50). The number of simulated variables were set at 50, 100, 200, 500, 1000 with corresponding sample sizes. (B) shows the box plots of estimated proteomic correlations for LEAP under larger sample sizes (10000, 20000, 50000, 100000, 200000) under the same proteomic explainability conditions as in (A). Subfigures (C) to (D) show the computational time (in seconds) required by each method across different ratios of the number of simulated variables to sample size (C) and larger sample sizes (D) under the same proteomic explainability conditions. (E) shows the box plots of estimated explainability for LEAP across different ratios of the number of simulated variables to sample size (N/M = 30 and N/M = 50) under high (set at 0.8) and low (set at 0.3) explainability conditions. (F) shows the estimated explainability for LEAP across larger sample sizes. Inside each box, the dash lines represent the true values, the solid lines indicate the median of the estimated values, the central boxes indicate the interquartile range (IQR), and the whiskers extend up to 1.5 times the IQR.

Although the computational time of LEAP increased with larger sample sizes, it remained manageable even at a sample size of 200,000 (Fig. 4C and Fig. 4D). Under these conditions, LEAP also provided unbiased estimates of proteomic explainability (Fig. 4E and Fig. 4F).

Additionally, we evaluated the standard errors of LEAP obtained from block jackknifing, which aligned well with the observed standard deviation values (Table S1). We also investigated the impact of residual correlations between two traits (ranging from 0.1 to 0.9). LEAP demonstrated robustness in proteomic correlation estimation across various settings of true residual correlations, maintaining accuracy under different scenarios of proteomic explainability (Fig. S1).

### Proteomic correlations among 50 traits in UKBB

To illustrate the effectiveness of LEAP in a large-scale cohort, we performed proteomic correlation analysis using LEAP based on the 1,461 proteins generated from the Olink Explore platform. The data were derived from plasma samples of 43,509 white British individuals in the UKB-PPP[9, 31]. We selected 50 traits encompassing anthropometric, behavioral, biochemical, and disease-related traits from the UKBB (Fig. 5A and Table S2). Out of the 1,225 pairwise combinations of these traits, 585 pairs exhibited significant correlations at the proteomic level after False Discovery Rate (FDR) correction. The study identified several clusters that could be classified into anthropometrics (e.g., weight, hip circumstance, waist circumstance, body mass index (BMI)), blood pressure (e.g., diastolic blood pressure, systolic blood pressure and pulse rate), kidney function (e.g., cystatin C, urea and creatinine), and red blood cell (e.g., red blood cell count, haemoglobin concentration and haematocrit percentage). The aforementioned traits exhibited high proteomic correlations in each cluster.

**Fig. 5.**
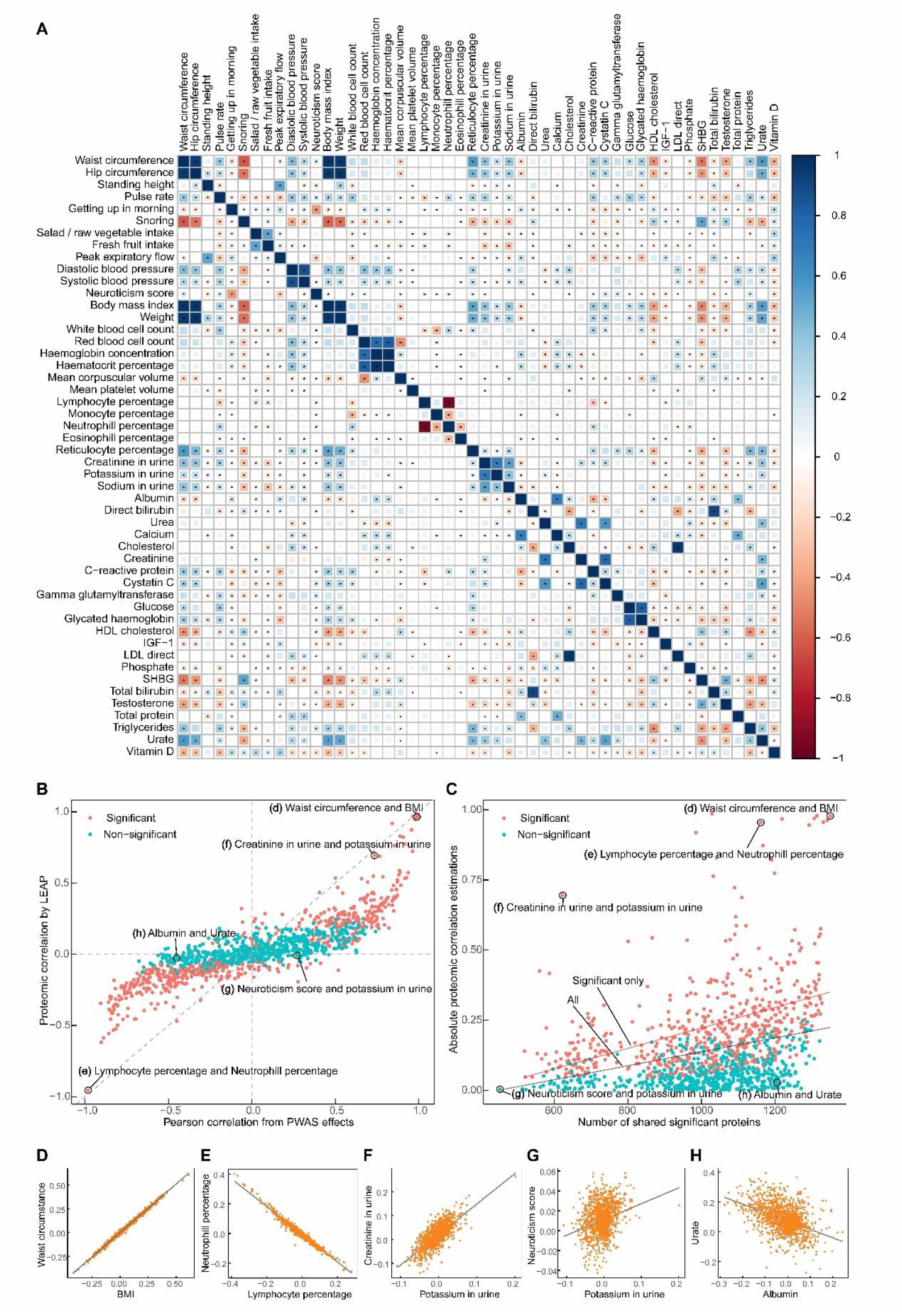
The proteomic correlation estimates in the UKBB. (A) The heat map plot displays the proteomic correlation estimates using LEAP among 50 traits at the protein level. The areas of the squares represent the absolute value of corresponding proteomic correlations. After FDR correction for 1,225 tests at a 5% significance level, proteomic correlation estimates that are significantly different from zero (star) are shown. (B) The scatterplot displays the association between proteomic correlation estimates (x-axis) and Pearson correlations of PWAS effects (y-axis) in the UKBB. The color represents the significance of the proteomic correlation estimates of the 1225 trait pairs. The grey cross and diagonal dashed line represent identity. (C) The scatterplot displays the association between the number of shared significant proteins after FDR correction in PWAS (x-axis) and the absolute proteomic correlation estimates (y-axis) are shown. The two solid lines represent the fitted lines of all dots and significant dots respectively. (D-H) Five scatterplots of five examples show the PWAS effects of 10 traits in the UKBB. The black line represents the Pearson correlation of each pair of traits.

It is noteworthy that the proteomic correlations were positively correlated but not quite equal with the measured effect detected by PWAS (Fig. 5B) and the numbers of shared significant proteins between each pair of traits (Fig. 5C). Traits with both high PWAS effects correlations and a substantial proportion of shared significant proteins, tended to exhibit high correlations at the proteomic level. For example, there was a proteomic correlation of 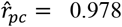 (*P* = 4.429 × 10^−250^) between waist circumference and BMI, both important measures of central obesity [32], with a PWAS effects correlation *r*_*coef*_ = 0.996 (Fig. 5D) and 1350 (out of 1461 proteins) shared significant proteins. Similarly, the proteomic correlation between lymphocyte percentage and neutrophil percentage was -0.955 (*P* = 5.059 × 10^−38^), with a PWAS effects correlation *r*_*coef*_ = -0.982 (Fig. 5E) and 1161 shared significant proteins.

Traits with high PWAS effects and a moderate proportion of shared significant proteins could also be significantly correlated at the proteomic level. For instance, the correlation between creatinine and potassium in the urine had 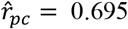 (*P* = 1.813 × 10^−8^), *r*_*coef*_ = 0.735 (Fig. 5F), and 622 shared significant proteins. This finding aligns with the previous studies implicating a biological association between Urine Chemistries [33]. However, traits with low PWAS effects correlation might exhibit non-significant proteomic correlations, irrespective of the proportions of shared significant proteins. Examples include neuroticism score and potassium in urine (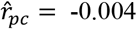, *P* = 0.828), and albumin and urate (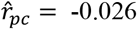, *P* = 0.691) (Fig. 5G and Fig. 5H). A comprehensive table of all 1,225 correlations among 50 traits is provided in Table S3.

Fitting a naive LMM on such a large size cohort of UKBB is computationally very demanding[30]. In this case, we utilized the modified LMM to estimate proteomic correlations with the UKBB. The point estimations of LMM were consistent with the results by LEAP (the Pearson correlation was approximately equal to 1, *P* < 2 × 10^−16^), indicating that LEAP is comparable to LMM (Fig. 6A and Fig. S2).

**Fig. 6.**
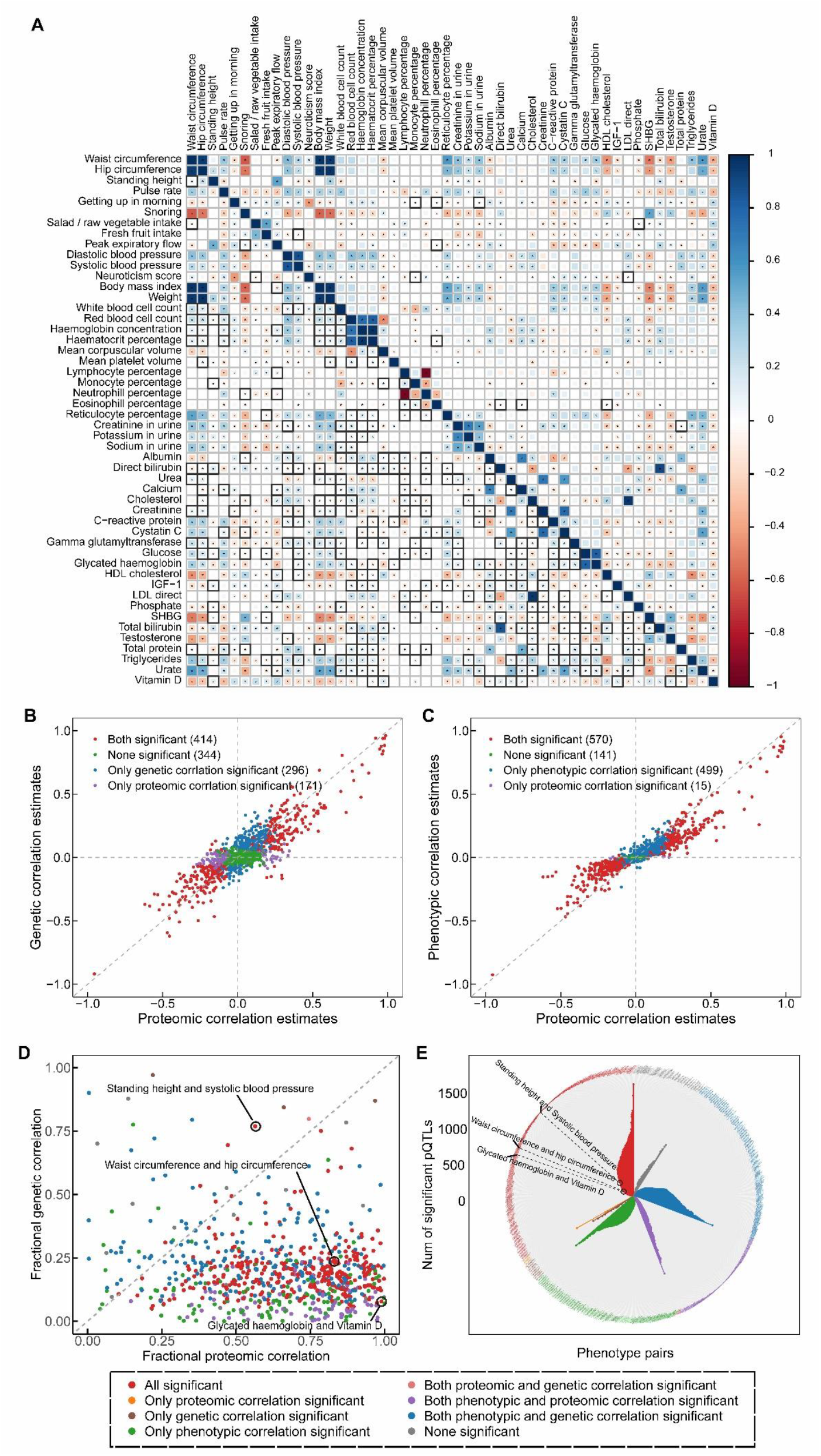
Real data application results in the UKBB. (A) Proteomic correlation estimates using LEAP and LMM among 50 traits at the protein level. Upper triangle: LEAP estimates; lower triangle: LMM estimates. The areas of the squares represent the absolute value of corresponding proteomic correlations. After FDR correction for 1,225 tests at a 5% significance level, proteomic correlation estimates that were significantly different from zero in both methods (star) and only one method (star and black square) are shown. (B) The scatterplot displays the performance comparison of the proteomic correlation estimates using LEAP (x-axis) and phenotypic correlations using Spearman’s Rank Correlation analysis (y-axis) among the 50 traits. Data points are color-coded based on significance, with the significant number of trait pairs in brackets. The grey diagonal dashed line represents identity. (C) shows the comparison results between proteomic correlation estimates and genetic correlation estimates. (D) The scatterplot displays the contributions of proteomic and genetic correlations to phenotypic correlations. The fractional proteomic correlations (x-axis) and the fractional genetic correlations (y-axis) of the 1225 trait pairs were calculated in all participants. Notable phenotype pairs (e.g., standing height and systolic blood pressure; waist circumference and hip circumference) are labeled. (E) The circular plot shows the significant QTLs (Quantitative Trait Loci) for each trait pair. Data points are color-coded based on significance categories. Notable phenotype pairs are labeled around the circle.

### Comparison of proteomic correlation with genetic and phenotypic correlations among 50 traits in UKBB

#### Comparison of proteomic correlation with genetic correlations among 50 traits in UKBB

The comparison of proteomic and genetic correlations among 50 traits in the UKBB revealed insights into the overall similarities and differences in phenotypic correlations at the proteomic and genetic levels. We utilized LDSC[7, 34] to estimate genetic correlations for the 50 traits in the UKBB, resulting in 1,225 genetic correlations. These were then compared to proteomic correlations estimated by LEAP. For most pairs of traits, the genetic correlation estimates closely matched the proteomic ones (the Pearson correlation is 0.855, *P* < 2 × 10^−16^), indicating a strong overall agreement. However, notable discrepancy was observed between the two correlation types. Among these, 414 pairs of traits were significantly correlated at both proteomic and genetic levels, and an additional 296 pairs of traits were observed as significant only for genetic correlations, compared to 171 pairs that were significant only for proteomic correlations (Fig. 6B and Fig. S3). Our findings aligned with existing biological knowledge in many instances. For example, standing height, a trait heavily influenced by genetics[35], showed genetic correlations with most anthropometric and behavioral traits. However, at the proteomic level, it lacked correlations with waist circumference, BMI, and several behavioral traits, suggesting distinct protein regulatory pathways. On the other hand, neuroticism score was not found to be genetically correlated with biochemical traits such as C-reactive protein, as suggested by the previous studies[36]. However, at the proteomic level, a significant correlation was observed between neuroticism score and C-reactive protein, hinting at potential shared regulatory pathways at the protein level. This implies that the mechanisms influencing the genetic and proteomic aspects of traits may diverge, providing valuable insights into the biological associations between mental disorders and inflammatory markers[37].

#### Comparison of proteomic and phenotypic correlations among 50 traits in UKBB

We estimated the Spearman correlation between each pair of the 50 traits to represent the phenotypic correlations, identifying 1069 pairs of significantly correlated traits after FDR correction. The majority of proteomic-correlated traits exhibited significant correlations at the phenotypic level, mirroring observations for genetically correlated traits (Fig. 6C and Fig. S4). Additionally, there were 15 pairs of traits with significant proteomic correlations but non-significant phenotypic correlations. For instance, the diastolic blood pressure and neuroticism score displayed a significant phenotypic correlation but lacked significant proteomic or genetic correlations. This suggests the existence of potential biological mechanisms between these two traits that warrant further investigations[38, 39].

#### Contribution of genetics and proteomics to phenotypic correlation

Different omics may have varying contributions to phenotypic correlations. To quantitatively compare the relative contributions of genetics and proteomics to phenotypic correlations, we introduced a relative statistical measure, 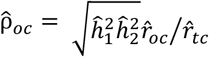 to express the contributions of different omics correlations to phenotypic correlation (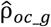 for genetics and 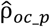 for proteomics), which is derived from the work of Searle[3] and Rheenen et al.[4] (See details in Methods and Table S4).

As shown in Fig. 6D, our results suggest that more trait pairs exhibit a higher proteomic contribution to phenotypic correlations compared to genetic contributions. For instance, the correlation between waist circumference and hip circumference is influenced more by their shared proteomic architectures 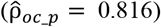 than by genetic factors 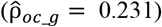. This finding is consistent with the observation that the proteomic correlation of these traits is greater than the genetic correlation 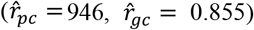. Similar patterns are observed for glycated hemoglobin and vitamin D, where high proteomic correlation and contribution 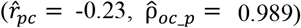 contrast with low genetic correlations and contributions 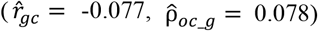.

#### pQTLs contribution to genetic and proteomic correlations

Protein quantitative trait loci (pQTL) play a crucial role in unraveling the genetic basis of protein-mediated genetic and disease associations[12]. Consequently, we investigated the presence and extent of pQTLs contributing to genetic and proteomic correlations (Fig. 6E). Leveraging the pQTL study conducted by Sun et al[11], we identified 1,363 proteins associated with 9,716 pQTLs (Table S5). For trait pairs exhibiting significant phenotypic, proteomic, and genetic correlations, we observed a substantial number of shared pQTLs among the traits. For instance, Cholesterol and LDL direct exhibited the highest number of shared pQTLs, with 1,550 pQTLs, and showed strong and significant correlations across all phenotypic, proteomic, and genetic levels. Notable examples include 73 significant pQTLs for the waist circumference and hip circumference pair, 100 pQTLs for standing height and systolic blood pressure pair, and 80 pQTLs for glycated haemoglobin and vitamin D pair.

These shared pQTLs between two traits suggest that the genetic contribution to phenotypic correlations may be mediated through proteins. Conversely, trait pairs with non-significant proteomic correlations displayed few significant pQTLs. For instance, no shared pQTL was found for the pulse rate and systolic blood pressure, and only one significant pQTL was identified for the neuroticism score and C-reactive protein (Table 2). This indicates the potential existence of alternative biological mechanisms through which genes influence these traits[40].

**Table 2.** Five examples of the PQTL analysis of the shared genetic variants and shared proteins.

| Pheno1 | Pheno2 | Shared<br>proteins | Shared<br>SNPs | PQTLs - shared<br>proteins | PQTLs - shared<br>SNPs | PQTLs - shared SNPs and<br>proteins |
| --- | --- | --- | --- | --- | --- | --- |
| Waist circumference | Hip circumference | 1293 | 4657 | 8985 | 110 | 73 |
| Standing height | Systolic blood<br>pressure | 1036 | 1687 | 6948 | 210 | 100 |
| Glycated<br>haemoglobin | Vitamin D | 1184 | 119 | 8221 | 158 | 80 |
| Pulse rate | Systolic blood<br>pressure | 1184 | 144 | 8051 | 0 | 0 |
| Neuroticism score | C-reactive protein | 708 | 577 | 5219 | 1 | 1 |

#### Differences in phenotypic, proteomic, and genetic correlations between male and female

The UKB-PPP dataset comprises 43,509 participants, with 23,456 females and 20,053 males (including self-identified genders and those identified through sex chromosome checks using genetic data). Our analysis revealed significant differences in most of the 50 traits between the two gender groups at the phenotypic level (Table S6). Consequently, we stratified the estimation of phenotypic, proteomic, and genetic correlations among these traits based on gender.

Overall, the results of gender-stratified correlations align closely with those obtained without gender stratification. However, substantial differences between males and females emerge at different levels of correlation. At the phenotypic level, among the 1,225 pairs of traits, 866 pairs of phenotypic correlations were detected in both gender groups, 134 pairs were specific to females, and 127 pairs were significant only in males (Fig. S5A, Fig. S5C, and Table S7). For instance, the correlation between testosterone and triglycerides was significant only in males (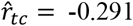, *P* < 2 × 10^−16^), but not significant in females (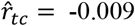, *P* = 0.197).

At the proteomic level, out of the 1,225 pairs of traits, 425 proteomic-correlated pairs were observed in both females and males, 196 pairs were exclusive to females, and 84 pairs were unique to males (Fig. 7A and Fig. S5D). Similarly, at the genetic level, 433 genetic-correlated pairs were identified in both female and male groups, 199 pairs were found only in females, and 107 pairs were specific to males (Fig. S5B and Fig. S5E). Consequently, there are more significant trait pairs in females compared to males. Numerous variations in proteomic and genetic correlations were identified between genders. For instance, in females, waist circumference and direct bilirubin exhibited significant correlations at both the proteomic level (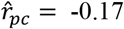, *P* = 9.629 × 10^−4^) and genetic level (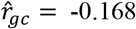, *P* = 2.443 × 10^−5^). However, in males, there was non-significance for both proteomic correlation (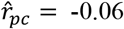, *P* = 0.117) and genetic correlation (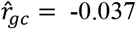, *P* = 0.32).

**Fig. 7.**
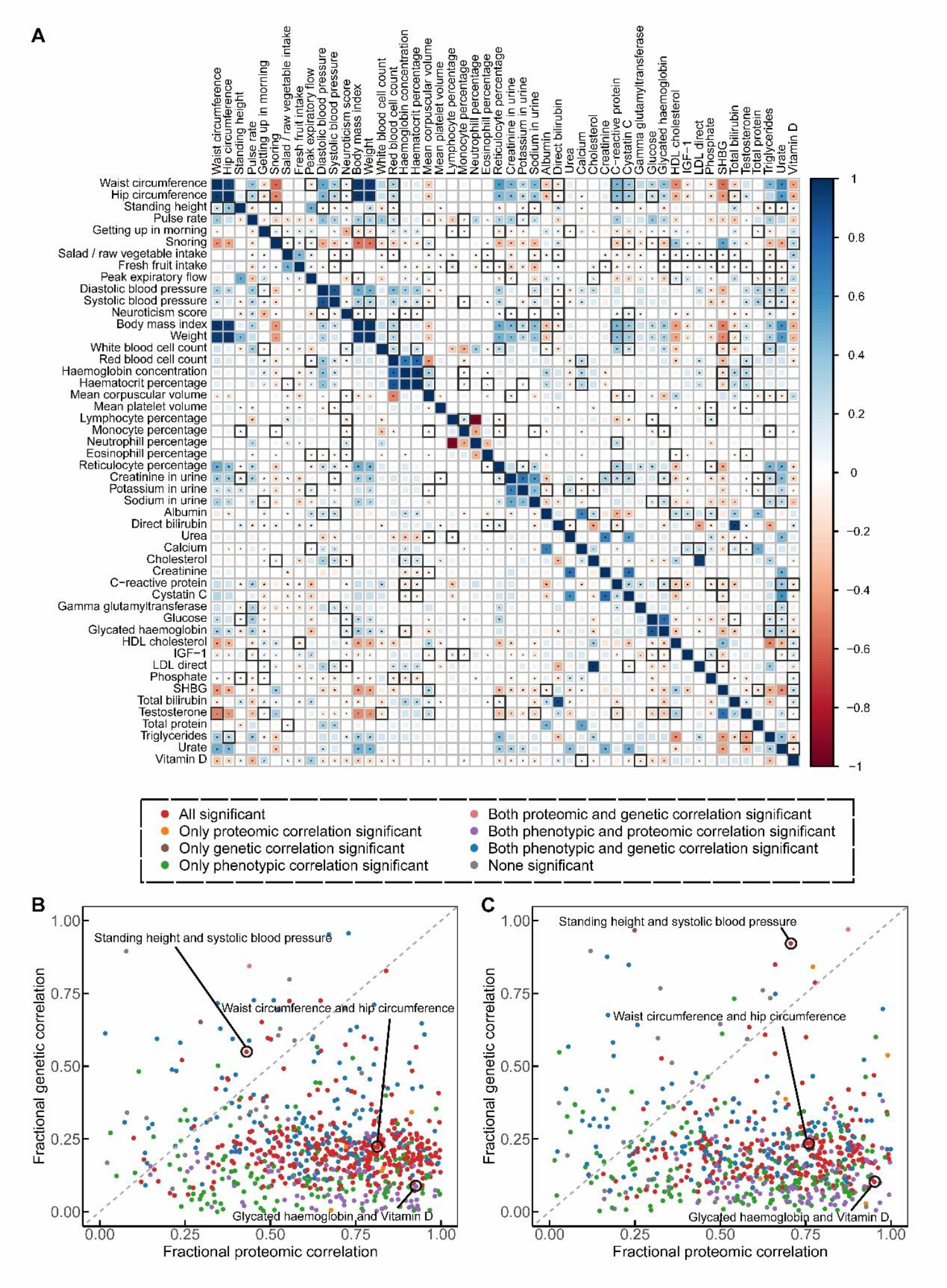
Gender differences in real data application results in the UKBB. (A) Gender differences in proteomic correlations among 50 traits. Upper triangle: females; lower triangle: males. The areas of the squares represent the absolute value of corresponding proteomic correlations. After FDR correction for 1,225 tests at a 5% significance level, proteomic correlation estimates that were significantly different from zero in both gender groups (star) and only one group (star and black square) are shown. (B) and (C) shows the gender differences in the contributions of proteomic and genetic correlations to phenotypic correlations. (B) shows the fractional proteomic correlations (x-axis) and the fractional genetic correlations (y-axis) of the 1225 trait pairs in females and (C) in males. The color represents the significance of the proteomic correlation estimates of the 1225 trait pairs. Data points are color-coded based on significance categories. Notable phenotype pairs are labeled around the circle.

Furthermore, a notable discrepancy exists in the proteomic and genetic contributions between male and female groups (Fig. 7B and Fig. 7C). For instance, both proteomic and genetic contributions to the phenotypic correlation between standing height and systolic blood pressure are lower in females (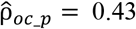 for proteomic contributions and 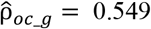 for genetic contributions) compared to males (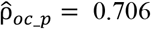 for proteomic contributions and 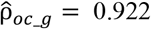 for genetic contributions).

## Discussion

In this study, we develop a novel likelihood-based approach, LEAP, for detecting and quantifying the shared proteomic architectures between two phenotypes. The LEAP method is based on estimating variance and covariance components from PWAS summary statistics, adjusting for covariance matrixes of proteomic data. Under the likelihood estimation framework, LEAP yields unbiased estimations of proteomic correlation across a wide range of scenarios and offers improved computational efficiency compared to existing methods. We demonstrated the advantages of LEAP based on extensive simulations and empirical cases from the UKBB. Additionally, we integrated phenotypic correlations with proteomic and genetic correlations to elucidate the shared biological underpinnings of these traits, highlighting how proteomics and genetics contribute differently to phenotypic correlations.

Genetic data are assumed to be independent across distance in genetic correlation methods. In contrast, proteins are influenced by both environmental risk factors and genetic variations[41], often resulting in high correlations even across different chromosomes[23]. This contrasts with genetic correlation approaches like LDSC[7, 19] and HDL[21], which assume independence across long-distance genome blocks and utilize partial covariance information to improve computational performance. The methodological assumptions behind these approaches are not suitable for estimating proteomic correlations. LEAP, in contrast, estimates proteomic correlations using orthogonal vectors and the covariance matrix of proteomic data separately, relaxing data architecture assumptions. Consequently, LEAP can be widely extended to estimating any shared non-genetic architectures among phenotypes, including epigenetics, transcriptomics, proteomics, and metabolomics.

For empirical cases, fitting a naive LMM model to a large-scale cohort like UKBB is computationally intensive, requiring either *O*(*MN*^2^) or *O*(*M*^2^*N*) runtime[42] and ∼O(*N*^2^) memory[6], where *N* is the sample size and *M* is the number of proteins. To address this in the context of proteomic correlations in UKBB, we computed the eigen-decomposition of the protein data to fit the LMM model, reducing computational time to ∼O(*N*)[6, 42, 43]. However, this process may lose efficacy in estimating proteomic variance (Fig. 2). Different from LMM, LEAP can handle large proteomic covariance matrix by performing eigen-decomposition on the covariance matrix of the proteomic data, utilizing the top eigenvalues and eigenvectors. For large sample sizes (*N*) but low-dimensional (*M*) omics data, the computational time is induced to ∼O(*M*), minimally affected by larger sample sizes. For high-dimensional proteomic data, only top eigenvalues and eigenvectors are passed to the model, mitigating the impact of high dimensionality. Thus, LEAP offers substantial computational efficiency gains over LMM for large sample sizes and high-dimensional markers (Supplementary Note 1.3).

Although LEAP requires a larger sample size to achieve fully unbiased estimations of proteomic explainability (Fig. 3), it can still be an estimator for quantifying the proportion of specific proteomic explainability for a trait within a population. Simulation studies demonstrated that proteomic correlation estimates by LEAP were unbiased under various conditions, regardless of sample size. This robustness is attributed to the ratio of proteomic correlation, akin to genetic correlation estimations using LDSC and HDL, where biases in both the numerator and denominator align and thus cancel each other out[7, 19, 21]. Given these considerations, we mainly focused on applying LEAP to estimating proteomic correlations.

We primarily modeled quantitative traits in the theoretical derivation and simulations of LEAP. For binary traits, treating them as continuous variables allows LEAP to estimate proteomic correlations, which can be justified by recognizing the linear model as a first-order Taylor approximation to generalized linear models like logistic regression[44]. It is also striking to note that we focused on modeling individual-level data with LEAP. However, LEAP can be easily extended to use summary statistics, as detailed in the Supplementary Notes and implemented in the same software. Briefly, the summary statistics version of LEAP requires marginal effect size estimates of the proteins on traits, their standard errors, and the proteomic correlation matrix from homogeneous samples. We simulated individual-level data and calculated the corresponding summary statistics for LEAP. Simulations showed that results using summary statistics were consistent with those from individual-level data-based LEAP (Fig. S6).

## Conclusions

In conclusion, LEAP stands out as a highly accurate and efficient method for estimating proteomic correlations within the realm of complex human traits. Leveraging the advantages of proteomic data architectures, LEAP can seamlessly extend to various non-genetic datasets, facilitating cross-omics correlation estimates. It accommodates both individual-level and summary-level data modeling, enabling the integration and alignment of multi-omics data from diverse sources. This establishes a versatile computational approach for efficiently estimating non-genetic correlations, facilitating the analysis of multiple layers influencing the formation and correlation of complex traits.

## Methods

### Modeling and estimation of proteomic correlation

Suppose there is a cohort for two traits with sample sizes *N*. The dimension of the proteomic data is *M* in the cohort. X denotes a *N* × *M* matrix of the proteomic data after the linear orthogonal transformation and is scaled to mean zero and variance one. Suppose the confounding environment factors are absent and the quantitative traits are affected by X through the multiple linear models,

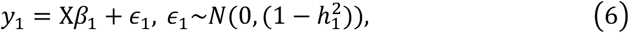

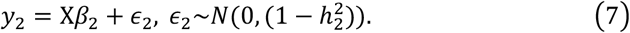

The *β*_*i*_-vector denotes the proteomic effects from X to *y*_*i*_,

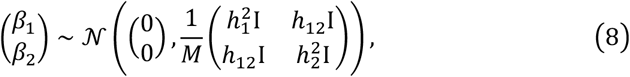

and residuals

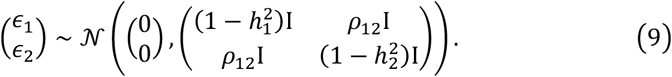

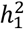 and 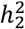 are the proteomic explainability. *h*_12_ is the proteomic covariance between two traits. *ρ*_12_ represents the covariance of residuals between the two traits. X, *β*, and *ϵ* are assumed to be independent of each other.

For phenotype *i*, the estimated marginal effect of X_*j*_ on *y*_*i*_ is

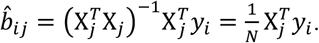

Then the covariance matrices are given as

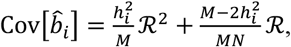

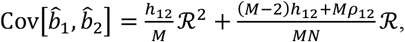

where ℛ is the *M* × *M* covariance matrix of the omics data. Let

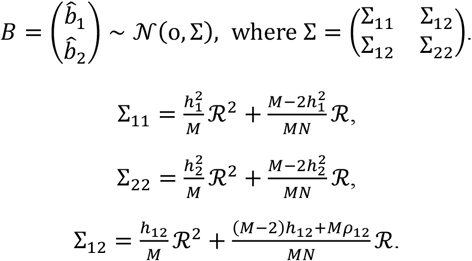

Noting the conditional distribution 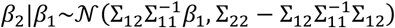, we can obtain the conditional log-likelihood

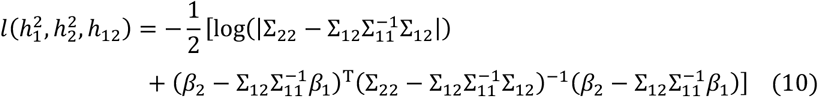

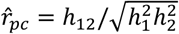 can be estimated by maximizing the conditional log-likelihood function to simplify the computation after eigen-decomposing the covariance matrix ℛ (see Supplementary Note 1.1-1.4 for more details).

### Simulations

We conducted a series of simulations to evaluate the performance of our method, LEAP, and compare it with the existing approach, LMM, as implemented in the *hglm* R package[45]. For LEAP, we performed eigen-decomposition on the matrix to transform the simulated proteomic data orthogonally. We then passed the eigenvalues and eigenvectors to LEAP to estimate the proteomic correlations. The standard error was computed using block-jackknifing (Supplementary Note 1.4), and the *P*-value was calculated using a two-sided Wald test with 1 degree of freedom. For LMM, we modeled the proteins as random correlated variables and treated the effect sizes as random effects, following previous genetic analysis studies[43, 46]. Considering the computational burden of LMM, we passed the eigenvalues and eigenvectors of the simulated protein data matrix to LMM to obtain the random effects on each trait. To distinguish between the two models, we refer to them as raw LMM and modified LMM, respectively. Proteomic correlations were then analyzed using the Pearson correlation coefficients between the random effects of each pair of traits. T-tests were used to compute the *P*-value for LMM.

We simulated *M* independent proteins from the multivariate normal distribution, X∼*N*(0, *I*). The true effect sizes *β*_1_ and *β*_2_ of X on the two traits were drawn from a bivariate normal distribution as formula (8). In addition, we simulated the two residual errors from another bivariate normal distribution as formula (9). Subsequently, the two quantitative traits, *y*_1_ and *y*_2_, were generated according to formulas (6) and (7). In the simulation, we first examined a baseline simulation setting where we set *N* = 3,000, *M* = 50, *h*_12_ = 0.5, 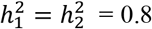, and *ρ*_12_= 0.1. Building on the baseline setting, we varied one parameter at a time to assess the influence of different parameters: the sample size *N* ranged from 1,000 to 5,000, the number of simulated proteins *M* ranged from 50 to 300, the true proteomic explainability of two traits 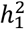 and 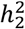 set at 0.8, and the true proteomic correlation *r*_*pc*_ varied from -0.9 to 0.9. We compared both the accuracy (difference between the simulated true value and estimated value) and efficiency (the average computational time for one simulation on a single 2.5 GHz Intel Core i7) among LEAP, raw LMM, and modified LMM for estimating the proteomic correlation of two traits and the proteomic explainability of each trait.

We further varied *N* from 1,000 to 20,000, *M* from 50 to 1,000, *r*_*pc*_ from -0.9 to 0.9, and *ρ*_12_ from 0.1 to 0.9 to assess the accuracy of LEAP, comparing with the modified LMM. Additionally, we tested the ratio of sample size to the number of simulated variables (N/M = 30 and N/M = 50) and larger sample sizes (N ranging from 10,000 to 200,000) to evaluate the robustness of LEAP. The proteomic explainability was configurated at three levels: both high 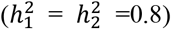, one high and one low (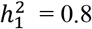 and 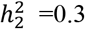), and both low 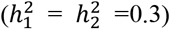 for two traits.

### Proteomic correlation estimates with UKBB

We estimated proteomic correlations of 50 traits using Olink Explore data from the UK Biobank (UKBB) Plasma Proteome Project (PPP), based on samples from 43,509 participants of White British ancestry[31]. Initially, 1463 unique proteins per participant were released in phase I. For this study, we analyzed 1461 proteins, excluding two proteins with a missing rate greater than 0.5. The remaining missing values were imputed using the mean value within the cohort. The 50 selected traits included anthropometric, behavioral, biochemical, and disease-related phenotypes from the UKBB. We adjusted for covariates such as age, sex, and assessment canters for each trait. Both LEAP and LMM were used to estimate the proteomic correlations. Previous studies have shown that LMM provides approximately unbiased cross-omics correlations using PC-based regularization of the simulated data matrix[47]. Considering the computational burden of LMM, we passed the eigenvalues and eigenvectors of the protein data matrix to LMM and used the modified LMM for proteomic correlation estimates.

### Genetic correlation estimates with UKBB

We estimated the genetic correlations of the same 50 traits in the UKBB using LDSC[7]. The UKBB GWAS summary statistics were obtained from the Pan-UKBB project, which provides GWAS summary statistics for over seven thousand phenotypes across six continental ancestry groups (available at https://pan.ukbb.broadinstitute.org/). We used the GWAS results from 420,531 European-ancestry individuals, updated in March 2023 by the Pan-UKBB team. After applying quality control (QC) filter, which included biallelic SNPs with INFO scores > 0.8, MAF > 1%, and exclusion of variants in the major histocompatibility complex region, we retained 1,096,114 autosomal variants from the imputed UKBB dataset. Covariates including age, sex, age × sex, age squared, age squared × sex, and the first 10 PCs of the genotype matrix were adjusted for each GWAS analysis. The linkage disequilibrium (LD) score files, specific to European ancestry, were also obtained from Pan-UKBB project. These files were computed on the QCed variants using a window size of 1 MB. We then estimated the genetic correlations using the *ldsc_rg* function implemented in the R package “*ldscr*”[34].

### Phenotypic correlation estimates with UKBB

We used Spearman’s Rank Correlation test to detect the phenotypic correlations among the 1,225 pairwise combinations of the 50 traits. Covariates such as age, sex, and sampling assessment center were adjusted for each trait.

### The contribution estimates of proteomic and genetic correlations to phenotypic correlations

As described by Searle[3] and Rheenen et al.[4], for standardized traits, the relationship between phenotypic correlation (*r*_*tc*_) and genetic correlation (*r*_*gc*_) is given by:

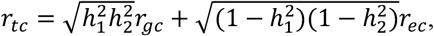

where 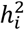 represents the explainability of each trait and *r*_*ec*_ is the additional correlation between residual factors. We can calculate the proportions of phenotypic correlations due to genetics using the fraction:

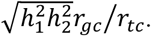

Conceptually, the genetic correlation can be replaced by the proteomic correlation with the corresponding proteomic explainability 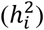 and the correlation between residual factors (*r*_*ec*_). We define the fraction parameter 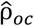 as:

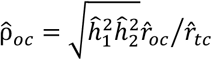

to indicate the contributions of different omics correlations (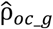 for genetics and 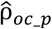 for proteomics) to the phenotypic interactions.

### PQTL analysis of shared genetic variants and shared proteins

We obtained comprehensive pQTL mappings from the UKBB based on the study by Sun et al[11]. The pQTL analyses were conducted on participants of European ancestry. Of the 1,461 proteins analyzed in this study, 1,363 proteins with 9,716 significant genetic associations were identified. For the 50 traits in the UKBB, we selected the shared significant proteins (*P* < 0.05 after Bonferroni correction) and the shared significant SNPs (P < 5×10^−8^) of each pair of traits. We then filtered and counted the significant genetic associations among these shared proteins and SNPs, including both *cis* and *trans* pQTLs.

## Ethics approval and consent to participate

This study was conducted using the UK Biobank Resource under application number 77803. UK Biobank has received ethical approval from the North West Multi-Centre Research Ethics Committee (MREC), which covers the ethical approval for the core UK Biobank study (REC reference: 11/NW/0382). All participants provided written informed consent to participate in the study, and all procedures followed were in accordance with institutional guidelines and regulations.

## Consent for publication

UK Biobank has obtained consent from all participants for their anonymized data to be used for scientific research and publication. This study was conducted using data from UK Biobank under application number 77803.

## Availability of data and materials

The data used in the present study are available from UK Biobank under restricted access. Data were used under license and are thus not publicly available. Access to the UKBB data can be requested through a standard protocol (https://www.ukbiobank.ac.uk/register-apply/). All data supporting the findings described in this manuscript are available within the article and the Supplementary Information. The UKBB GWAS summary statistics from the Pan-UKBB project were obtained from https://pan.ukbb.broadinstitute.org/.

Most operations are carried out by the software R 4.2.0 from CRAN (http://cran.r-project.org/). Code for LEAP is implemented in the R package *LEAP*, which is available in the Supplementary files (LEAP_0.1.0.tar.gz, which can be installed in R). LMM is performed by the R package *hglm* which is available at https://CRAN.R-project.org/package=hglm. LDSC is performed by the R package *ldscr* available at https://github.com/mglev1n/ldscr.

## Authors’ contributions

X.S., M.Z., S.W., and L.J. conceived the idea and designed the research; X.S., S.Y., and M.Z. evaluated the computational approaches. X.S., S.Y., Q.P., S.W., and X.Z performed simulations and real-data analysis. X.S. and Y.Q. developed the software. R.S. and G.Z. downloaded and preprocessed the real data. X.S., M.Z., Q.P., and L.J. wrote the manuscript. All authors reviewed and approved the final manuscript.

## Acknowledgements

We thank the UK Biobank resource, approved under application 77803, for the individual-level protein and clinical data.

