## Supplementary Information for "High-efficiency likelihood inference of shared proteomic architectures across 50 complex human traits"

### Authors and Affiliations

Xiaoru Sun[1, 2, #], Sizhe Yang[3, 4, 5, #], Qianqian Peng[6, #], Xuan Zhang[1], Yijia Qian[7], Renliang Sun[6], Guoqing Zhang[6], Sijia Wang[6, 8, *], Li Jin[1, 3, *], Menghan Zhang[1, 4, 5, 9, *]

[1] State Key Laboratory of Genetic Engineering, Human Phenome Institute, Fudan University, Shanghai, China

[2] Department of Medical Data, School of Public Health, Cheeloo College of Medicine, Shandong University, Jinan, China

[3] Collaborative Innovation Center for Genetics and Development, School of Life Sciences, Fudan University, Shanghai, China

[4] Ministry of Education Key Laboratory of Contemporary Anthropology, Department of Anthropology and Human Genetics, School of Life Sciences, Fudan University, Shanghai, China

[5] Research Institute of Intelligent Complex Sciences, Fudan University, Shanghai, China

[6] Key Laboratory of Computational Biology, Shanghai Institute of Nutrition and Health, University of Chinese Academy of Sciences, Chinese Academy of Sciences, Shanghai, China

[7] School of Software, Fudan University, Shanghai, China

[8] Center for Excellence in Animal Evolution and Genetics, Chinese Academy of Sciences, Kunming, China

[9] Lead contact

[#] These authors contributed equally.

[*] Corresponding authors: Menghan Zhang, Li Jin, Sijia Wang

### 1 Supplementary Note for the method LEAP

#### 1.1 Estimating the explainability of one trait using the likelihood method

Inspired by the development of the genetic correlation technique applying to GWAS summary statistics, such as LDSC and HDL[1-3], we developed LEAP to estimate the variance/covariance components based on the marginal association test statistics (which can be obtained from the proteome-wide association study) corrected for covariance matrixes of proteomic data. Suppose there is a cohort for two traits with sample sizes $n$. The dimension of proteomic data $Z$ is $m$ in the cohort. $y_1 = \{y_{1i}\}$ is a $n \times 1$ vector for trait 1; $y_2 = \{y_{2i}\}$ is a $n \times 1$ vector for trait 2. $Z$ is a $n \times m$ matrix of the proteomic data. Without loss of generality, $Z$ is scaled to mean zero and variance one. Suppose the confounding environmental factors are absent and we model the two quantitative traits $y_i$ ($i$ = 1, 2 for two traits respectively) as affected by $Z$ from the equations respectively,

$$y_i = \mathrm{Z}\alpha_i + e_i. \quad (1)$$

We assume all three variables on the right side of the above equation (1) are random. $\alpha_i$ is a $m$-vector of proteomic effect sizes on the $i$-th trait. $e_i$ is an $n$-vector of residual error and follows a matrix normal distribution $MN_{n,m}(0, \sigma^2 I)$. $\mathrm{X}$ is a $n \times m$ matrix of the proteomic data after the linear orthogonal transformation, i.e. $\mathrm{X} = ZV$, where $V$ is the right singular matrix of $Z$ ($Z = U\Lambda_1 V^T$). Without loss of generality, $\mathrm{X}$ is also scaled to mean zero and variance one. Then we can model the two quantitative traits $y_i$ as affected by $\mathrm{X}$ from the equation

$$y_i = \mathrm{X}\beta_i + \epsilon_i. \quad (2)$$

According to equations (1) and (2), we can obtain the effects $\beta_i = V^T\alpha_i$ and $cor(\beta_1,\ \beta_2) = cor(V^T\alpha_1,\ V^T\alpha_2) = cor(\alpha_1,\ \alpha_2)$. Thus, the proteomic correlation can be estimated via the effect size $\beta_i$. Based on the ordinary least squares estimation, $\beta_i = (X^T X)^{-1} X^T y_i$. Thus

$$\mathrm{E}(\beta_i) = \mathrm{E}[\mathrm{E}(\beta_i|X)] = E[E((X^T X)^{-1} X^T y_i|X)] = E[E(X^T)E(y_i|X)] = 0.$$

Then the two effect sizes $\beta_i$ follow the joint distribution

$$\begin{pmatrix}\beta_1\\ \beta_2\end{pmatrix} \sim \mathcal{N}\left(\begin{pmatrix}0\\ 0\end{pmatrix}, \frac{1}{m}\begin{pmatrix}h_1^2\mathrm{I} & h_{12}\mathrm{I}\\ h_{12}\mathrm{I} & h_2^2\mathrm{I}\end{pmatrix}\right),$$

and residuals

$$\begin{pmatrix}\epsilon_1\\ \epsilon_2\end{pmatrix} \sim \mathcal{N}\left(\begin{pmatrix}0\\ 0\end{pmatrix}, \begin{pmatrix}(1-h_1^2)\mathrm{I} & \rho_{12}\mathrm{I}\\ \rho_{12}\mathrm{I} & (1-h_2^2)\mathrm{I}\end{pmatrix}\right).$$

$h_1^2$ and $h_2^2$ are the proportion of specific proteomic variance in the population, which is here defined as the proteomic explainability. $h_{12}$ is the proteomic covariance between two traits. $\rho_{12}$ represents the covariance of residuals between the two traits. $\mathrm{X}, \beta,$ and $\epsilon$ are assumed to be independent of each other.

For phenotype *i*, the estimated marginal effect of $\mathrm{X}_j$ on $y_i$ is

$$\hat{b}_{ij} = \left(\mathrm{X}_j^T \mathrm{X}_j\right)^{-1} \mathrm{X}_j^T y_i = \frac{1}{n-1} \mathrm{X}_j^T y_i.$$

For one of the traits $y_1$,

$$y_1 = \mathrm{X}\beta_1 + \epsilon_1,\ \epsilon_1 \sim N(0, (1-h_1^2)).$$

The estimated marginal effect of $\mathrm{X}_j$ on $y_1$ is

$$\hat{b}_{1j} = \left(\mathrm{X}_j^T \mathrm{X}_j\right)^{-1} \mathrm{X}_j^T y_1 = \frac{1}{n-1} \mathrm{X}_j^T y_1.$$

$$
\begin{aligned}
\mathrm{E}\left[\hat{b}_{1j}\hat{b}_{1j'} \mid \mathrm{X}\right] &= \frac{1}{(n-1)^2}\mathrm{E}\left[\mathrm{X}_j^T y_1 y_1^T \mathrm{X}_{j'} \mid \mathrm{X}\right] \\
&= \frac{1}{(n-1)^2}\mathrm{E}\left[\mathrm{X}_j^T(\mathrm{X}\beta_1+\epsilon_1)(\mathrm{X}\beta_1+\epsilon_1)^T\mathrm{X}_{j'} \mid \mathrm{X}\right] \\
&= \frac{1}{(n-1)^2}\mathrm{E}\left[\mathrm{X}_j^T(\mathrm{X}\beta_1\beta_1^T\mathrm{X}^T+\epsilon_1\beta_1^T\mathrm{X}^T+\mathrm{X}\beta_1\epsilon_1^T+\epsilon_1\epsilon_1^T)\mathrm{X}_{j'} \mid \mathrm{X}\right] \\
&= \frac{1}{(n-1)^2}\mathrm{E}\left[\mathrm{X}_j^T(\mathrm{X}\beta_1\beta_1^T\mathrm{X}^T+\epsilon_1\epsilon_1^T)\mathrm{X}_{j'} \mid \mathrm{X}\right] \\
&= \frac{1}{(n-1)^2}\left(\mathrm{X}_j^T\mathrm{X}\mathrm{E}[\beta_1\beta_1^T \mid \mathrm{X}]\mathrm{X}^T\mathrm{X}_{j'}+\mathrm{X}_j^T\mathrm{E}[\epsilon_1\epsilon_1^T \mid \mathrm{X}]\mathrm{X}_{j'}\right) \\
&= \frac{1}{(n-1)^2}\left(\frac{h_1^2}{m}\mathrm{X}_j^T\mathrm{X}\mathrm{X}^T\mathrm{X}_{j'}+(1-h_1^2)\mathrm{X}_j^T\mathrm{X}_{j'}\right).
\end{aligned}
$$

Let $\hat{r}_{jk}=\mathrm{X}_j^T\mathrm{X}_k/(n-1)$, $\hat{r}_{j'k}=\mathrm{X}_{j'}^T\mathrm{X}_k/(n-1)$ and $\hat{r}_{jj'}=\mathrm{X}_j^T\mathrm{X}_{j'}/(n-1)$, then

$$
\mathrm{E}\left[\hat{b}_{1j}\hat{b}_{1j'} \mid \mathrm{X}\right]=\frac{h_1^2}{m}\sum_{k=1}^{m}\hat{r}_{jk}\hat{r}_{j'k}+\frac{1-h_1^2}{n-1}\hat{r}_{jj'}.
$$

Take expectation over X, we have

$$
\mathrm{E}\left[\hat{b}_{1j}\hat{b}_{1j'}\right]=\mathrm{E}\left[\mathrm{E}\left[\hat{b}_{1j}\hat{b}_{1j'} \mid \mathrm{X}\right]\right]=\frac{h_1^2}{m}\sum_{k=1}^{m}\mathrm{E}\left(\hat{r}_{jk}\hat{r}_{j'k}\right)+\frac{1-h_1^2}{n-1}\mathrm{E}\left(\hat{r}_{jj'}\right)
$$

By the law of large numbers, $\mathrm{E}\left[\hat{r}_{jj'}\right]=r_{jj'}$. For the expected value of $\hat{r}_{jk}\hat{r}_{j'k}$,

$$
\mathrm{E}\left[\hat{r}_{jk}\hat{r}_{j'k}\right]=\mathrm{E}\left[\hat{r}_{jk}\right]\mathrm{E}\left[\hat{r}_{j'k}\right]+\mathrm{Cov}\left[\hat{r}_{jk},\hat{r}_{j'k}\right]=r_{jk}r_{j'k}+\mathrm{Cov}\left[\hat{r}_{jk},\hat{r}_{j'k}\right].
$$

According to Pearson and Filon[4],

$$
\mathrm{Cov}\left[\hat{r}_{jk},\hat{r}_{j'k}\right]=\frac{1}{N-1}\left[r_{jj'}\left(1-r_{jk}^2-r_{j'k}^2\right)-\frac{1}{2}r_{jk}r_{j'k}\left(1-r_{jk}^2-r_{j'k}^2-r_{jj'}^2\right)\right].
$$

Because X is linear orthogonal transformed, $r_{jk}=0$ if $j\neq k$.

$$
\sum_{k=1}^{m}\mathrm{Cov}\left[\hat{r}_{jk},\hat{r}_{j'k}\right]=\frac{1}{n-1}\left[r_{jj'}\left(m-r_{jj}^2-r_{j'j'}^2\right)\right]=\frac{m-2}{n-1}r_{jj'}.
$$

Thus,

$$
\sum_{k=1}^{m}\mathrm{E}\left[\hat{r}_{jk}\hat{r}_{j'k}\right]=\sum_{k=1}^{m}r_{jk}r_{j'k}+\frac{m-2}{n-1}r_{jj'}=r_{jj'}^2+\frac{m-2}{n-1}r_{jj'}.
$$

Then,

$$\mathrm{E}\big[\hat{b}_{1j}\hat{b}_{1j'}\big] = \frac{h_1^2}{m}\sum_{k=1}^{m}\mathrm{E}\big(\hat{r}_{jk}\hat{r}_{j'k}\big) + \frac{1-h_1^2}{n-1}\mathrm{E}\big(\hat{r}_{jj'}\big) = \frac{h_1^2}{m}\Big(r_{jj'}^2 + \frac{m-2}{n-1}r_{jj'}\Big) + \frac{1-h_1^2}{n-1}r_{jj'}$$

$$= \frac{h_1^2}{m}r_{jj'}^2 + \frac{(m-2)h_1^2 + m\big(1-h_1^2\big)}{m(n-1)}r_{jj'} = \frac{h_1^2}{m}r_{jj'}^2 + \frac{m-2h_1^2}{m(n-1)}r_{jj'}.$$

Similarly, for the trait $y_2$, we can obtain $\mathrm{E}\big[\hat{b}_{2j}\hat{b}_{2j'}\big] = \frac{h_2^2}{m}r_{jj'}^2 + \frac{m-2h_2^2}{m(n-1)}r_{jj'}$.

### 1.2 Estimating proteomic correlations using the likelihood method

Considering two traits, the expected product of $\hat{b}_{1j}$ and $\hat{b}_{2j'}$ given X

$$\begin{aligned}
\mathrm{E}\big[\hat{b}_{1j}\hat{b}_{2j'} \mid \mathrm{X}\big] &= \frac{1}{(n-1)^2}\mathrm{E}\big[\mathrm{X}_j^T y_1 y_2^T \mathrm{X}_{j'} \mid \mathrm{X}\big] \\
&= \frac{1}{(n-1)^2}\mathrm{E}\big[\mathrm{X}_j^T(\mathrm{X}\beta_1 + \epsilon_1)(\mathrm{X}\beta_2 + \epsilon_2)^T\mathrm{X}_{j'} \mid \mathrm{X}\big] \\
&= \frac{1}{(n-1)^2}\mathrm{E}\big[\mathrm{X}_j^T(\mathrm{X}\beta_1\beta_2^T\mathrm{X}^T + \epsilon_1\beta_2^T\mathrm{X}^T + \mathrm{X}\beta_1\epsilon_2^T + \epsilon_1\epsilon_2^T)\mathrm{X}_{j'} \mid \mathrm{X}\big] \\
&= \frac{1}{(n-1)^2}\mathrm{E}\big[\mathrm{X}_j^T(\mathrm{X}\beta_1\beta_2^T\mathrm{X}^T + \epsilon_1\epsilon_2^T)\mathrm{X}_{j'} \mid \mathrm{X}\big] \\
&= \frac{1}{(n-1)^2}\big(\mathrm{X}_j^T\mathrm{X}\mathrm{E}[\beta_1\beta_2^T \mid \mathrm{X}]\mathrm{X}^T\mathrm{X}_{j'} + \mathrm{X}_j^T\mathrm{E}[\epsilon_1\epsilon_2^T \mid \mathrm{X}]\mathrm{X}_{j'}\big) \\
&= \frac{1}{(n-1)^2}\Big(\frac{h_{12}}{m}\mathrm{X}_j^T\mathrm{X}\mathrm{X}^T\mathrm{X}_{j'} + \rho_{12}\mathrm{X}_j^T\mathrm{X}_{j'}\Big).
\end{aligned}$$

Let $\hat{r}_{jk} = \mathrm{X}_j^T\mathrm{X}_k/(n-1)$, $\hat{r}_{j'k} = \mathrm{X}_{j'}^T\mathrm{X}_k/(n-1)$ and $\hat{r}_{jj'} = \mathrm{X}_j^T\mathrm{X}_{j'}/(n-1)$, then

$$\mathrm{E}\big[\hat{b}_{1j}\hat{b}_{2j'} \mid \mathrm{X}\big] = \frac{h_{12}}{m}\sum_{k=1}^{m}\hat{r}_{jk}\hat{r}_{j'k} + \frac{n-1}{(n-1)^2}\rho_{12}\hat{r}_{jj'} = \frac{h_{12}}{m}\sum_{k=1}^{m}\hat{r}_{jk}\hat{r}_{j'k} + \frac{\rho_{12}}{n-1}\hat{r}_{jj'}.$$

Take expectation over X, we have

$$\mathrm{E}\big[\hat{b}_{1j}\hat{b}_{2j'}\big] = \mathrm{E}\Big[\mathrm{E}\big[\hat{b}_{1j}\hat{b}_{2j'} \mid \mathrm{X}\big]\Big] = \frac{h_{12}}{m}\sum_{k=1}^{m}\mathrm{E}\big[\hat{r}_{jk}\hat{r}_{j'k}\big] + \frac{\rho_{12}}{n-1}\mathrm{E}\big[\hat{r}_{jj'}\big]$$

$$= \frac{h_{12}}{m}\Big(r_{jj'}^2 + \frac{m-2}{n-1}r_{jj'}\Big) + \frac{\rho_{12}}{n-1}r_{jj'} = \frac{h_{12}}{m}r_{jj'}^2 + \frac{(m-2)h_{12} + m\rho_{12}}{m(n-1)}r_{jj'}.$$

Let the $m \times m$ matrix $\mathcal{R} = \{r_{jk}\}$ denotes the covariance matrix of $X$, where $r_{jk} =$

$\mathrm{E}[\mathrm{X_j X_k}]$. Let

$$B = \begin{pmatrix} \hat{b}_1 \\ \hat{b}_2 \end{pmatrix} \sim \mathcal{N}(\mathrm{o}, \Sigma), \text{ where } \Sigma = \begin{pmatrix} \Sigma_{11} & \Sigma_{12} \\ \Sigma_{12} & \Sigma_{22} \end{pmatrix}.$$

$$\Sigma_{11} = \frac{h_1^2}{m}\mathcal{R}^2 + \frac{m-2h_1^2}{m(n-1)}\mathcal{R},$$

$$\Sigma_{22} = \frac{h_2^2}{m}\mathcal{R}^2 + \frac{m-2h_2^2}{m(n-1)}\mathcal{R},$$

$$\Sigma_{12} = \frac{h_{12}}{m}\mathcal{R}^2 + \frac{(m-2)h_{12}+m\rho_{12}}{m(n-1)}\mathcal{R}.$$

$h_1^2, h_2^2, h_{12}$ can be estimated by maximizing the full joint log-likelihood function,

$$\max l(h_i^2) = \max\left\{-\frac{1}{2}\left[\log(|\Sigma_{ii}|) + \beta_i^{\mathrm{T}}\Sigma_{ii}^{-1}\beta_i\right]\right\},$$

$$\text{and } \max l(h_1^2, h_2^2, h_{12}) = \max\left\{-\frac{1}{2}\left[\log(|\Sigma|) + B^{\mathrm{T}}\Sigma^{-1}B\right]\right\}.$$

It is noted that the conditional distribution for $\beta_2$ given $\beta_1$ is

$$\beta_2|\beta_1 \sim \mathcal{N}(\Sigma_{12}\Sigma_{11}^{-1}\beta_1, \Sigma_{22} - \Sigma_{12}\Sigma_{11}^{-1}\Sigma_{12}).$$

This gives the conditional log-likelihood function

$$l(h_1^2, h_2^2, h_{12}) = -\frac{1}{2}[\log(|\Sigma_{22} - \Sigma_{12}\Sigma_{11}^{-1}\Sigma_{12}|)$$

$$+ (\beta_2 - \Sigma_{12}\Sigma_{11}^{-1}\beta_1)^{\mathrm{T}}(\Sigma_{22} - \Sigma_{12}\Sigma_{11}^{-1}\Sigma_{12})^{-1}(\beta_2 - \Sigma_{12}\Sigma_{11}^{-1}\beta_1)].$$

**1.3 Using eigen-decomposition to simplify computation**

As a real symmetric matrix, $\mathcal{R}$ can be decomposed as $\mathcal{R} = Q\Lambda Q^T$, where $Q$ is an orthogonal matrix whose columns are the eigenvectors of $\mathcal{R}$ (a unit matrix as $X$ is a orthogonally transformed matrix), and $\Lambda$ is the diagonal matrix whose entries are the eigenvalues of $\mathcal{R}$, $\Lambda = \begin{pmatrix} \lambda_1 & \cdots & 0 \\ \vdots & \ddots & \vdots \\ 0 & \cdots & \lambda_m \end{pmatrix}$. Then $\Sigma_{ii}$ and $\Sigma_{12}$ can be reformed to

$$\Sigma_{ii} = \frac{h_i^2}{m}\mathcal{R}^2 + \frac{m-2h_i^2}{m(n-1)}\mathcal{R} = Q\left(\frac{h_i^2}{m}\Lambda^2 + \frac{m-2h_i^2}{m(n-1)}\Lambda\right)Q^T,$$

$$\Sigma_{11}^{-1} = Q\left(\frac{h_1^2}{m}\Lambda^2 + \frac{m-2h_1^2}{m(n-1)}\Lambda\right)^{-1} Q^T,$$

$$\Sigma_{12} = \frac{h_{12}}{m}\mathcal{R}^2 + \frac{(m-2)h_{12}+m\rho_{12}}{m(n-1)}\mathcal{R} = Q\left(\frac{h_{12}}{m}\Lambda^2 + \frac{(m-2)h_{12}+m\rho_{12}}{m(n-1)}\Lambda\right)Q^T.$$

Then $l(h_i^2)$ can be transformed to

$$l(h_i^2) = -\frac{1}{2}\left[\sum_{j=1}^{m}\log\left(\frac{h_i^2}{m}\lambda_j^2 + \frac{m-2h_i^2}{m(n-1)}\lambda_j\right) + \beta_i^{\mathrm{T}} Q\left(\frac{h_1^2}{m}\Lambda^2 + \frac{m-2h_1^2}{m(n-1)}\Lambda\right)^{-1} Q^T\beta_i\right].$$

Denoting $u_i = Q^T\beta_i$ with entries $\{u_{ij}\}$, and denoting $\lambda_{ii,j}^{*} = \frac{h_i^2}{m}\lambda_j^2 + \frac{m-2h_i^2}{m(n-1)}\lambda_j$, we have

$$l(h_i^2) = -\frac{1}{2}\left[\sum_{j=1}^{m}\log\left(\frac{h_i^2}{m}\lambda_j^2 + \frac{m-2h_i^2}{m(n-1)}\lambda_j\right) + u_i^{\mathrm{T}}\left(\frac{h_1^2}{m}\Lambda^2 + \frac{m-2h_1^2}{m(n-1)}\Lambda\right)^{-1} u_i\right]$$

$$= -\frac{1}{2}\left[\sum_{j=1}^{m}\log\left(\lambda_{ii,j}^{*}\right) + \sum_{j=1}^{m}\frac{u_{ij}^2}{\lambda_{ii,j}^{*}}\right].$$

Similarity, denoting $\lambda_{12,j}^{*} = \frac{h_{12}}{m}\lambda_j^2 + \frac{(m-2)h_{12}+m\rho_{12}}{m(n-1)}\lambda_j$, we can obtain that

$$\Sigma_{22} - \Sigma_{12}\Sigma_{11}^{-1}\Sigma_{12} = Q\left[\left(\frac{h_2^2}{m}\Lambda^2 + \frac{m-2h_2^2}{m(n-1)}\Lambda\right) - \left(\frac{h_{12}}{m}\Lambda^2 + \frac{(m-2)h_{12}+m\rho_{12}}{m(n-1)}\Lambda\right)\left(\frac{h_1^2}{m}\Lambda^2 + \frac{m-2h_1^2}{m(n-1)}\Lambda\right)^{-1}\left(\frac{h_{12}}{m}\Lambda^2 + \frac{(m-2)h_{12}+m\rho_{12}}{m(n-1)}\Lambda\right)\right]Q^T,$$

then $\log(|\Sigma_{22} - \Sigma_{12}\Sigma_{11}^{-1}\Sigma_{12}|) = \sum_{j=1}^{m}\log\left(\lambda_{22,j}^{*} + \frac{\left(\lambda_{12,j}^{*}\right)^2}{\lambda_{11,j}^{*}}\right)$.

Let $u^{*} = Q^T[\beta_2 - \Sigma_{12}\Sigma_{11}^{-1}\beta_1]$, $l(h_1^2, h_2^2, h_{12})$ can be transformed to

$$l(h_1^2, h_2^2, h_{12}) = -\frac{1}{2}\left[\sum_{j=1}^{m}\log\left(\lambda_{22,j}^{*} + \frac{\left(\lambda_{12,j}^{*}\right)^2}{\lambda_{11,j}^{*}}\right) + \sum_{j=1}^{m}\frac{\left(u_j^{*}\right)^2}{\lambda_{22,j}^{*} - \frac{\left(\lambda_{12,j}^{*}\right)^2}{\lambda_{11,j}^{*}}}\right].$$

**1.4 Estimating standard variation of proteomic correlations**

Firstly, we randomly sampled 70% individuals (0.7×$m$) in the simulation dataset or in the real dataset each time to estimate the pseudo correlations ($\hat{r}_m'$) using LEAP. 200

replications ($K$) were performed. Then we could obtain the standard variation of proteomic correlations $Var(r_m) = \frac{0.7\times m}{m}\sum_{k=1}^{K}(\hat{r}_m' - \bar{r}_m')^2$ that was similar to the block- jackknife method and $\bar{r}_m' = mean(\hat{r}_m')$. The s.e. values from block jackknifing were consistent with the observed s.d. values (Supplementary Table 1)

#### 1.5 Using PWAS summary statistics to estimate proteomic correlations

To make LEAP applicable for summary-level data, we used the PWAS summary statistics (the marginal regression effect $\mu$) and the proteomic covariance matrix ($Z^T Z$) to estimate proteomic correlations and compared them with the estimates obtained from the individual-level data. Theoretically, the proteomic data $Z$ can be eigen-decomposed as $Z = U\Lambda_1 V^T$ and the proteomic covariance matrix can be eigen-decomposed as $Z^T Z/(n-1) = \mathrm{U}\Lambda_1 V^T \mathrm{V}\Lambda_1 U^T = \mathrm{V}\Lambda_1^2 V^T = \mathrm{V}\Lambda V^T$, where $\Lambda$ is the eigenvalue of $\mathcal{R}$ ($\mathcal{R} = Q\Lambda Q^T$) and $\Lambda = \Lambda_1^2/(n-1)$. The orthogonal matrix $Q = V^T V$. It is also noted that the orthogonally transformed proteomic data matrix $\mathrm{X} = ZV$.

For phenotype $i$, the originally estimated marginal effect of $Z$ on $y_i$ is

$$\hat{\mu} = \left(tr(Z^T Z)\right)^{-1} Z^T y_i = \left(tr(\mathrm{V}\Lambda V^T)\right)^{-1} V\Lambda_1 U^T y_i = \frac{1}{n-1} V\Lambda_1 U^T y_i.$$

The transformed marginal effect of $X$ on $y_i$ is

$$\hat{b} = \left(tr(X^T X)\right)^{-1} X^T y_i = \Lambda^{-1} V^T V \Lambda_1 U^T y_i = \Lambda_1^{-1} U^T y_i = \Lambda^{-1} V^T \hat{\mu}.$$

Then, the parameters $h_1^2, h_2^2, h_{12}$ in the full joint log-likelihood function can be estimated using the summary statistics as the same as using the individual-level data.

### 2 Supplementary Tables

**Supplementary Table 1: Simulations with different heritability groups for the standard variation of proteomic correlation estimations.**

| Heritability group | Method | True value | Estimate | s.d. | s.e. |
|---|---|---|---|---|---|
| **High** | LEAP | 0.5 | 0.53 | 0.02 | 0.02 |
| | LMM | 0.5 | 0.51 | 0.021 | 0.027 |
| **Low** | LEAP | 0.5 | 0.5 | 0.023 | 0.024 |
| | LMM | 0.5 | 0.46 | 0.025 | 0.028 |

- Sample size *n*: 10000
- Proteins *m*: 1000
- High $R^2$: 0.8
- Low $R^2$: 0.3
- Observed s.d.: sd of 200 simulations
- True s.e.: median of s.e. from 2000 jackknifes among 200 simulations

**Supplementary Table 2: The detailed descriptions of 3 covariates and 50 traits in the UKBB.**

[See the Excel file]

**Supplementary Table 3: 1225 proteomics correlations among 50 traits in the UKBB.**

[See the Excel file]

**Supplementary Table 4: The contribution of proteomic and genetic correlations to phenotypic correlations in the UKBB.**

[See the Excel file]

**Supplementary Table 5: PQTL analysis of the shared genetic variants and shared**

**proteins in the UKBB.**

[See the Excel file]

**Supplementary Table 6: The statistical tests of 50 traits with gender in the UKBB.**

[See the Excel file]

**Supplementary Table 7: 1225 proteomics correlations among 50 traits in different gender groups in the UKBB.**

[See the Excel file]

### 3 Supplementary Figures

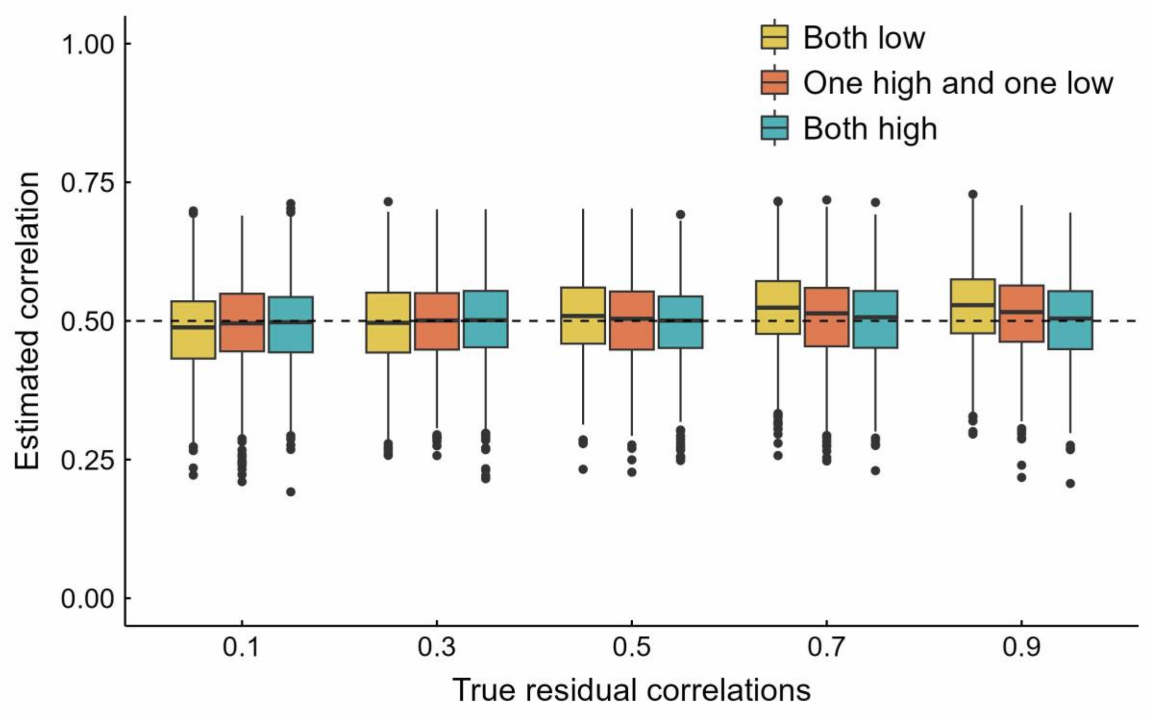


**Supplementary Figure 1. Proteomic correlation estimates using LEAP under different levels of residual correlation.** In each proteomic explainability group and residual correlation level, 1000 replicates were simulated. The true proteomic correlation was set at 0.5. The high proteomic explainability of both two traits was set at 0.8 and the low proteomic explainability at 0.3. 5000 of 100 independent variables were randomly generated as simulated proteins. Inside each box, the dashed lines represent the true values, the solid lines indicate the median of the estimated values, the central boxes indicate the interquartile range (IQR), and the whiskers extend up to 1.5 times the IQR.

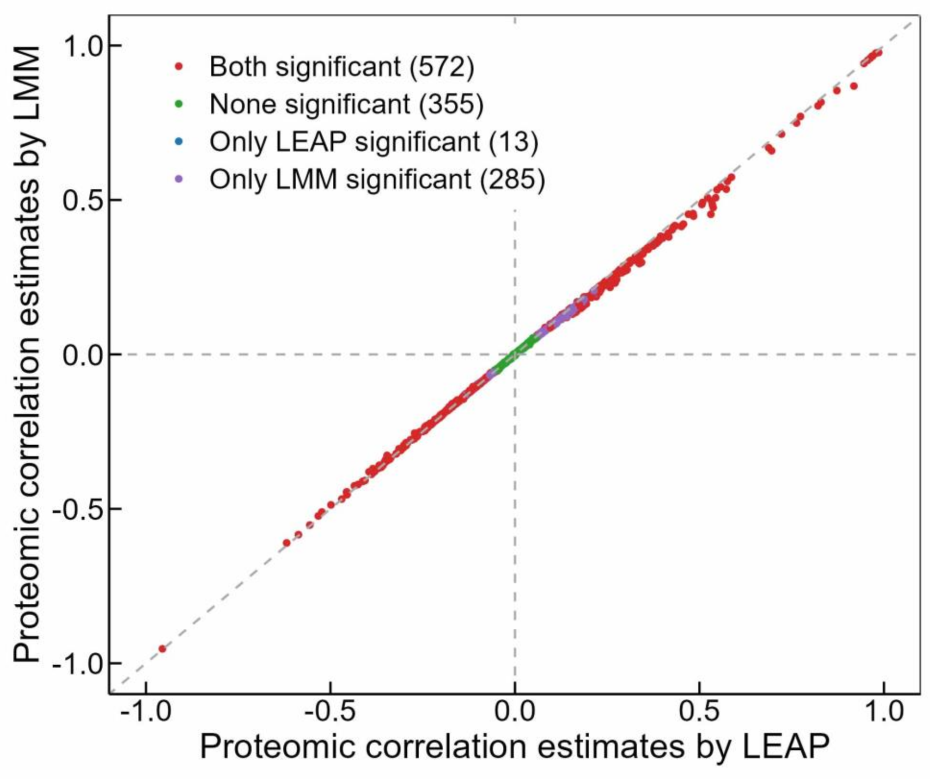


Supplementary Figure 2: Scatterplots to display the proteomic correlation estimates using LEAP (x-axis) and LMM (y-axis) with the UKBB cohort.

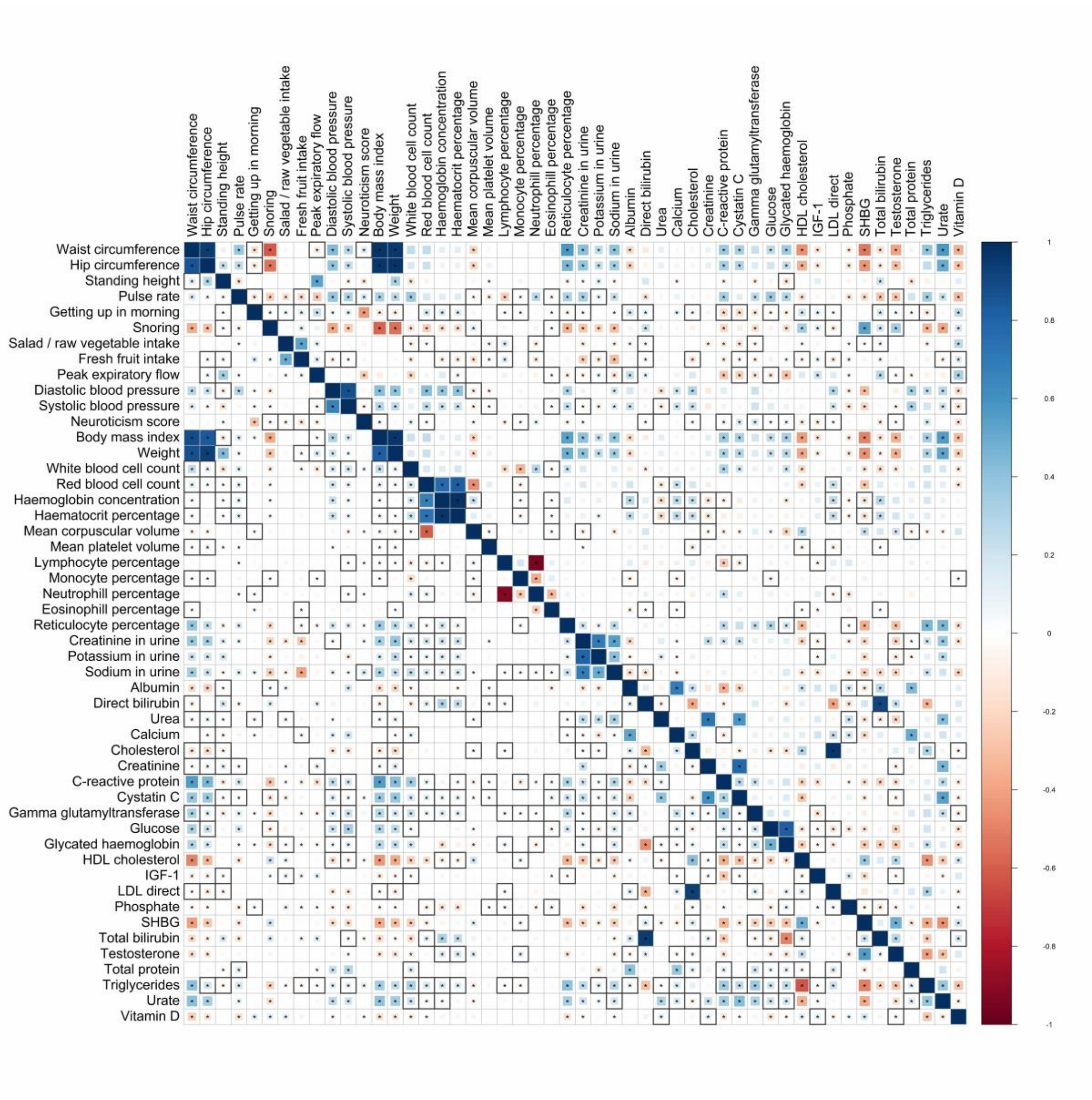


Supplementary Figure 3: Proteomic correlation estimates using LEAP (upper triangle) and genetic correlations using LDSC (lower triangle) among the 50 traits. The areas of the squares represent the absolute value of corresponding proteomic correlations. After FDR correction for 1,225 tests at a 5% significance level, both proteomic correlation and genetic correlation estimates that were significantly different from zero (star) and only one significant (star and black square) are shown.

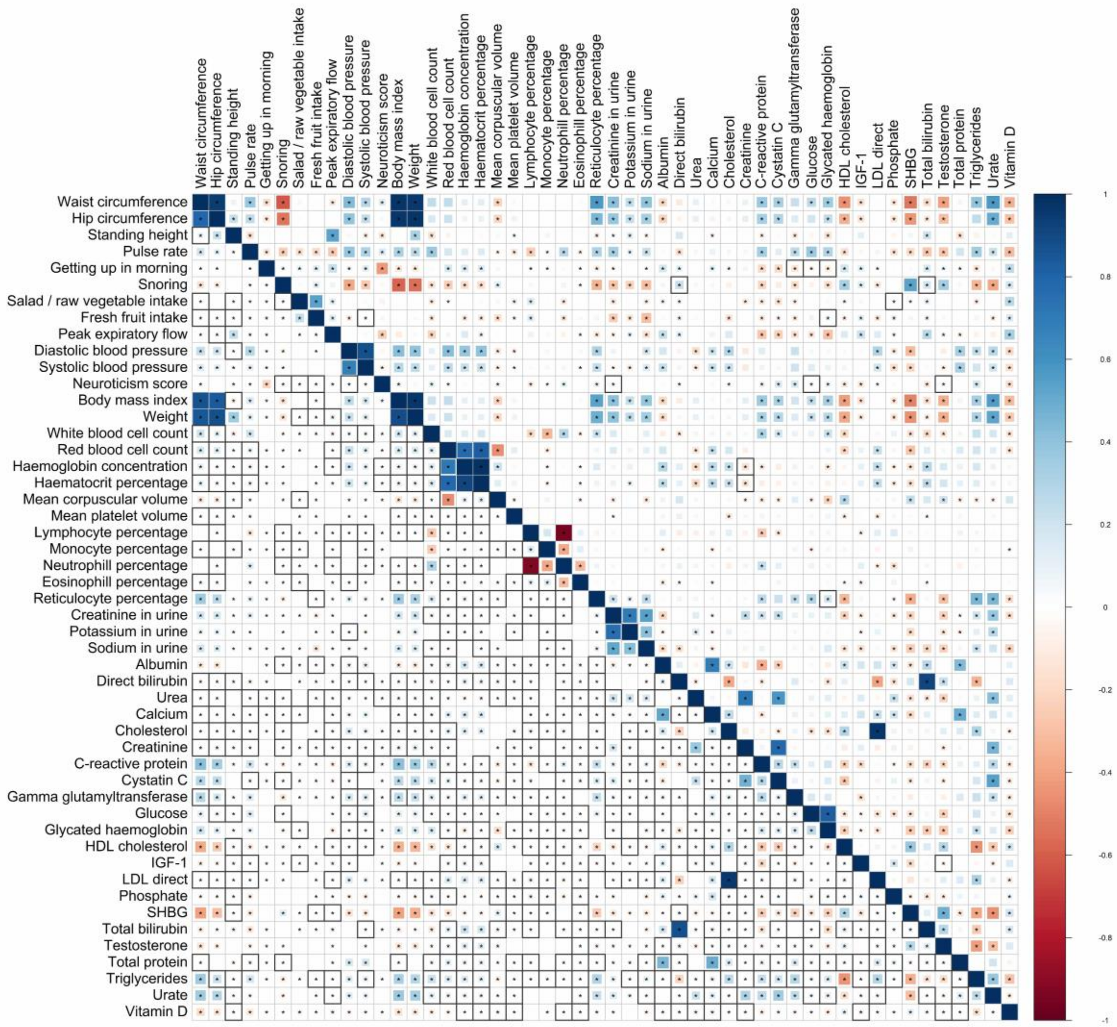


Supplementary Figure 4: Proteomic correlation estimates using LEAP (upper triangle) and phenotypic correlations using Spearman's Rank Correlation analysis (lower triangle) among the 50 traits. The areas of the squares represent the absolute value of corresponding proteomic correlations. After FDR correction for 1,225 tests at a 5% significance level, both proteomic correlation and phenotypic correlation estimates that were significantly different from zero (star) and only one significant (star and black square) are shown.

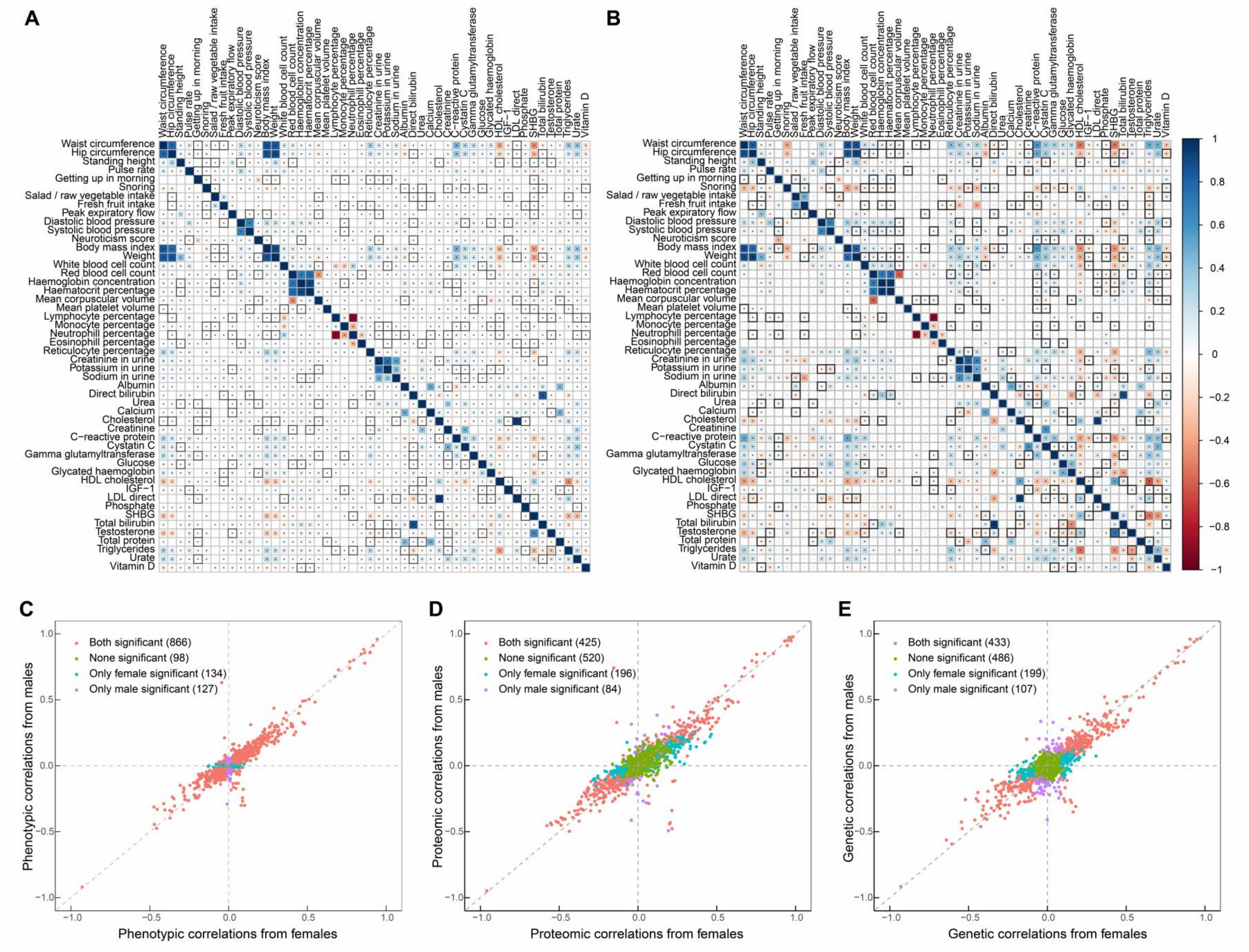
A
B
Waist circumference
Hip circumference
Standing height
Pulse rate
Getting up in morning
Snoring
Salad / raw vegetable intake
Fresh fruit intake
Peak expiratory flow
Diastolic blood pressure
Systolic blood pressure
Neuroticism score
Body mass index
Weight
White blood cell count
Red blood cell count
Haemoglobin concentration
Haematocrit percentage
Mean corpuscular volume
Mean platelet volume
Lymphocyte percentage
Monocyte percentage
Neutrophill percentage
Eosinophill percentage
Reticulocyte percentage
Creatinine in urine
Potassium in urine
Sodium in urine
Albumin
Direct bilirubin
Urea
Calcium
Cholesterol
Creatinine
C−reactive protein
Cystatin C
Gamma glutamyltransferase
Glucose
Glycated haemoglobin
HDL cholesterol
IGF−1
LDL direct
Phosphate
SHBG
Total bilirubin
Testosterone
Total protein
Triglycerides
Urate
Vitamin D
1
0.8
0.6
0.4
0.2
0
−0.2
−0.4
−0.6
−0.8
−1
C
Both significant (866)
None significant (98)
Only female significant (134)
Only male significant (127)
Phenotypic correlations from males
Phenotypic correlations from females
D
Both significant (425)
None significant (520)
Only female significant (196)
Only male significant (84)
Proteomic correlations from males
Proteomic correlations from females
E
Both significant (433)
None significant (486)
Only female significant (199)
Only male significant (107)
Genetic correlations from males
Genetic correlations from females
−1.0
−0.5
0.0
0.5
1.0

Supplementary Figure 5: Gender differences in estimating genetic correlations and phenotypic correlations in the UKBB. (A) Phenotypic correlations among 50 traits in the UKBB. Upper triangle: females; lower triangle: males. The areas of the squares represent the absolute value of corresponding phenotypic correlations. After FDR correction for 1,225 tests at a 5% significance level, phenotypic correlation estimates that were significantly different from zero in both gender groups (star) and only one group (star and black square) are shown. (B) Genetic correlations among 50 traits in the UKBB. Upper triangle: females; lower triangle: males. (C) Scatterplots to display the gender differences in estimating phenotypic correlations. (D) Scatterplots to display the gender differences in estimating proteomic correlations using LEAP. (E) Scatterplots to display the gender differences in estimating genetic correlations using LDSC.

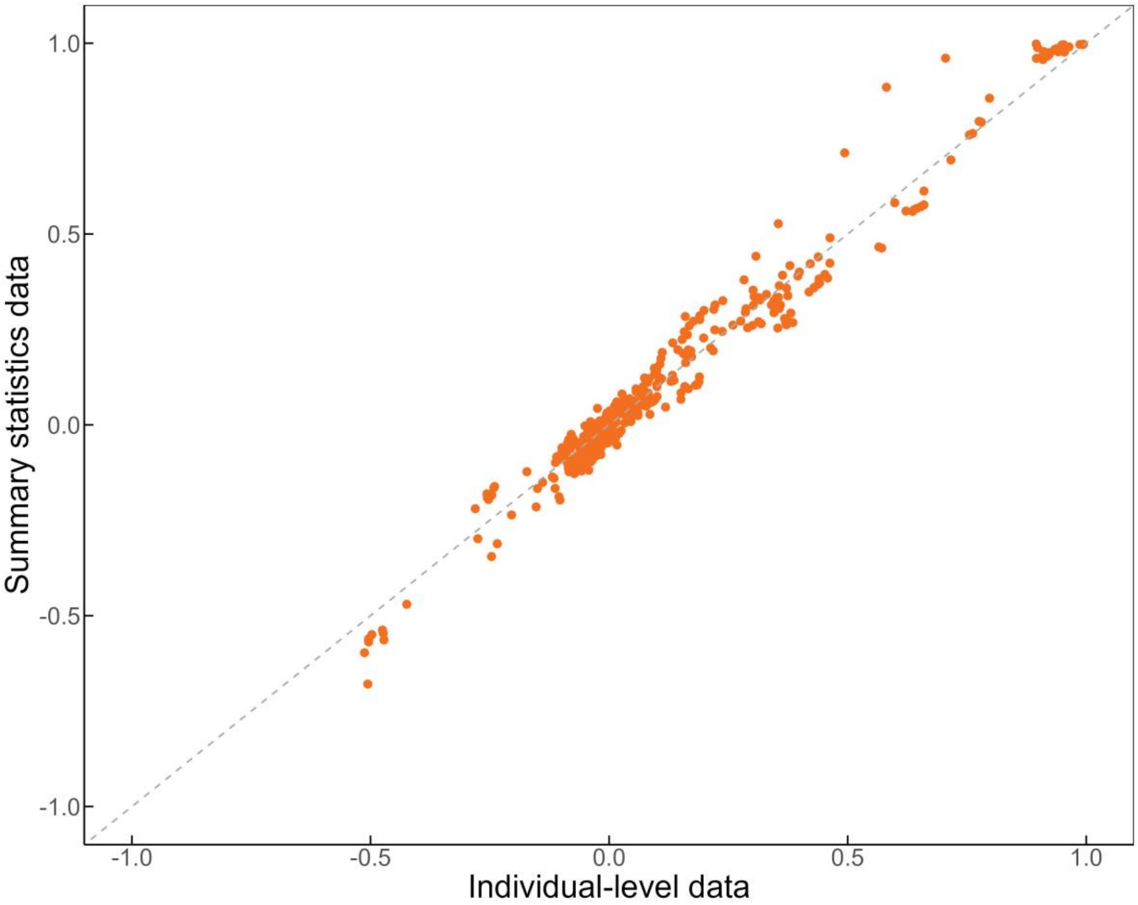


Supplementary Figure 6: Comparing proteomic correlation estimates using LEAP with individual-level data (x-axis) and summary statistics data (y-axis) among 30 simulated traits (435 pairs of traits).
